# Biomarker Variability Limits Individualized Amyloid Time Estimation in Alzheimer Disease

**DOI:** 10.64898/2026.07.08.737258

**Authors:** Julie K. Wisch, Ziqiao Jiao, Peter R. Millar, Nicole S. McKay, Aleksandra Beric, Wenjing Lin, Bryce Baker, Jennifer Stauber, Sam Preminger, Mathias Jucker, Nicolas R. Barthélemy, Jasmeer Chhatwal, Natalie S Ryan, Suzanne E. Schindler, John C. Morris, Carlos Cruchaga, Tammie L.S. Benzinger, Celeste M. Karch, Randall J. Bateman, Eric McDade, Jorge Llibre-Guerra, The Dominantly Inherited Alzheimer Network, The Alzheimer Disease Neuroimaging Initiative, Brian A. Gordon, Beau M. Ances, Laura Ibanez

## Abstract

**Objective:** Disease progression modeling (DPM) or “amyloid time” is increasingly used to stage Alzheimer disease (AD). DPM performance depends on within-individual heterogeneity in rates of pathological accumulation as well as test-retest reliability of the biomarker. The relative contributions of these variabilities have not been systematically assessed. This would be particularly relevant if extrapolations from DPM were to be used to make individual-level predictions for research, clinical trials, or potentially future clinical practice.

**Methods:** We conducted simulation studies incorporating empirically-derived noise properties from amyloid biomarkers to assess the contributions of inter- and intra-individual variability. Findings generalized in an autosomal dominant AD cohort with amyloid positron emission tomography (PET), cerebrospinal fluid (CSF), and plasma biomarkers and in a sporadic AD cohort with both amyloid PET and plasma biomarkers. We assessed group level DPM performance via mean average error (MAE) and root mean squared error (RMSE). At the individual level, we evaluated distinctness of distributions of biomarker levels associated with specific disease timings.

**Results:** Inter-individual variability was the dominant source of error in temporal estimates. Intra-individual variability reduced estimate stability. Optimal performance occurred in biomarkers with positive average accumulation rates where a subset of individuals had exceptionally high levels of accumulation. In research study data, amyloid PET outperformed CSF and plasma biomarkers.

**Interpretation:** DPM is fundamentally constrained by dynamic range, variability, and test–retest reliability of the biomarker of interest. Current DPM approaches are more robust at the group level, particularly when applied to biomarkers with more than 10-15% variability like fluid biomarkers.

**Funding:** National Institute on Aging, Alzheimer’s Association, German Center for Neurodegenerative Diseases, Raul Carrea Institute for Neurological Research, Japan Agency for Medical Research and Development, Korean Ministry of Health & Welfare and Ministry of Science and ICT, Spanish Institute of Health.

## INTRODUCTION

Disease Progression Modeling (DPM) reconstructs the temporal course of a disease by integrating short-term longitudinal observations across individuals^1^, enabling estimation of key disease milestones that cannot be observed within a single study. In Alzheimer’s Disease (AD), DPM has been used to estimate each individual’s temporal position relative to the onset of detectable amyloid accumulation (“amyloid time” or “amyloid chronicity”)^2–6^, a key initiating event in AD ^7^. Although these estimates are typically interpreted at the group level, extending DPM to the individual level could facilitate personalized disease staging, supporting precision medicine efforts. However, both inter- and intra-individual variability have the potential to distort disease timeline estimates. A study that explicitly evaluates their effects on DPM performance is necessary to evaluate the appropriateness of DPM for individual level predictions.

DPM was originally applied to amyloid PET^2,3,8,9^, but has recently been extended to fluid biomarkers.^6,10–12^ A critical gap in the DPM literature is that the fundamental modeling assumptions of uniform pathological accumulation for a given baseline biomarker level and negligible within-biomarker variability have not been rigorously evaluated. These assumptions likely vary by biomarker modality and should be evaluated as expansion of this technique to more biomarkers is considered.

Recent efforts to apply DPM to plasma p-tau181 resulted in substantial uncertainty particularly at the upper bounds of the DPM-derived estimates due to high measurement variability^13^. Plasma p-tau217 assays, which have shown a stronger continuous relationship with amyloid PET^14,15^, may outperform plasma p-tau181, but this requires further validation. Further, most published DPM studies rely on relatively short follow-up periods, meaning that model performance is evaluated primarily among individuals who convert near the time of observation. Consequently, accuracy across the full preclinical disease trajectory remains largely unknown. Although one previous study combined simulations with application to an AD cohort,^8^ it did not evaluate the effects of inter- or intra-individual variability impact on model performance.

Here, we systematically investigated how intra- and inter-individual variability affect DPM performance using both simulation and observational data. Simulations spanning the full preclinical disease duration were constructed using empirically derived biomarker characteristics. We then evaluated DPM performance in the Dominantly Inherited Alzheimer Network (DIAN), selected for its breadth of biomarker characterization,^7,8,15–20^ and the Alzheimer’s Disease Neuroimaging Initiative (ADNI) as an independent validation cohort representing sporadic AD (sAD). Resampling analyses were performed throughout to assess model stability and generalizability.

## METHODS

This study applied DPM using a parallel approach with simulation and observational research datasets. Simulation studies systematically characterized model performance across a wide range of realistic conditions, and the observational analyses were conducted in the DIAN and ADNI cohorts to evaluate generalizability of simulation findings to collected biomarker data (Figure 1).

**Figure 1.**
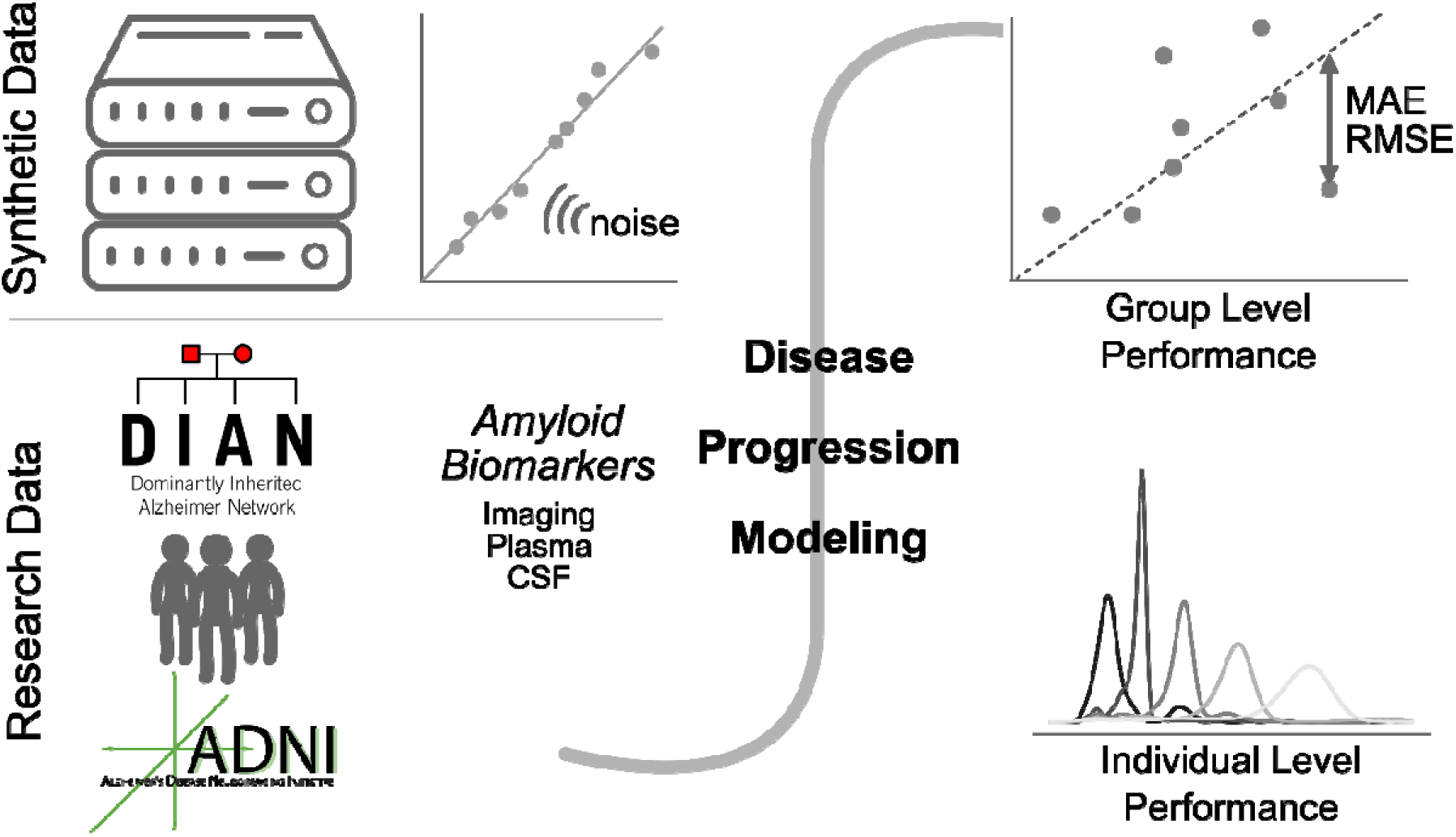
Study Design. Disease Progression Modeling (DPM) was assessed using a parallel approach for synthetic and research data (that had amyloid biomarkers available). DPM was applied to each synthetic dataset generated first in the absence of intra-individual noise and then at escalating levels (from 3% to 20%) and also to the longitudinal research datasets. Performance was evaluated at the group level using mean absolute error (MAE) and root mean square error (RMSE). Performance at individual level was evaluated by assessing the distinctness of distributions of biomarker levels associated with specific disease timings.

### Data Sources

#### Simulation Study

We generated synthetic longitudinal biomarker data based on empirical values from research cohorts for 300 individuals. The series of visits for each individual were randomly sampled from a temporal distribution mirroring the sampling frequency of the DIAN study (mean=2.2 years, standard deviation=1.06 years). Each was assigned a baseline biomarker level sampled probabilistically to enrich for cases in which conversion from biomarker-negative to biomarker-positive status occurred during follow-up.

Annualized rate of change (ARC) was simulated using a skew-normal distribution to reflect the observed distribution of biomarker trajectories. This type of distribution is defined by three key features: the central tendency (ξ), skew (α), and spread (ω). Inter-individual variability was induced by systematically varying these parameters across plausible ranges derived from measured biomarker data (Supplemental Figure 1, 2). Intra-individual variability was introduced by adding Gaussian noise ranging from 3% to 20%, consistent with reported test–retest variability in PET and fluid biomarkers^16–19^.

#### Observational Study - DIAN

The DIAN Observational Study is a multi-site international research consortium that recruits individuals from families that carry an AD causing mutation in *APP, PSEN1*, or *PSEN2*. Written consent was obtained for all participants, and study protocols were approved by local institutional review boards of all DIAN sites.

DIAN participants (347 mutation carriers and 212 non-carriers; Supplemental Table 1) who had completed longitudinal amyloid PET, CSF and/or plasma biomarker measurements were included in this study. Non-mutation carrying DIAN participants were only used for norming when biomarker values were converted to Z scores. We assessed CSF p-tau181 measured using immunoprecipitation mass spectrometry (IP-MS; %pTau181), Lumipulse, and NULISA; CSF p-tau217 quantified with IP-MS (%p-tau217) and NULISA. CSF brain derived [BD] p-tau217 and p-tau181 were also available for a subset of individuals measured using NULISA. In plasma, p-tau181 was measured with Simoa, IP-MS, and NULISA; p-tau217 was quantified using IP-MS or NULISA (Supplemental Table 1). Batch correction was applied as necessary using bridging samples when available, and COMBAT when they were not^20^. PET and biofluid processing details are available in the methods supplement.

#### Observational Study - ADNI

The Alzheimer’s Disease Neuroimaging Initiative (ADNI) is a multi-site longitudinal research study designed to develop and validate biomarkers for sAD. Values included in this study were collected under ADNI4. Written informed consent was obtained from all participants, and study protocols were approved by the institutional review boards at participating sites.

We included 108 ADNI participants (Supplemental Table 2) who had longitudinal amyloid PET and plasma p-tau217/Aβ42 measured using Lumipulse Fujerubio^21^. We only included participants with paired values to keep the number of samples in ADNI to be of similar magnitude to that of available data in DIAN. Methods for biomarker collection have been previously published^22,23^. We applied long-COMBAT to harmonize amyloid PET across sites^24^.

#### Observational Study - Defining Key Characteristics

We first defined the threshold for biomarker positivity for each fluid biomarker, which is considered “Time 0” in DPM. We performed cutpoint analyses on each fluid biomarker that optimized classification accuracy for a threshold of 18 Centiloids (CL)^25^ using the Youden index (Supplemental Figure 2). We established a threshold for reliable accumulation for each biomarker, to ensure that only participants likely to exhibit biologically meaningful pathological change were included^4,26^. We performed individual-by-individual linear regression on available longitudinal data to obtain the ARC estimate for each participant. Finally, we calculated the central tendency, skew and spread for the ARC distribution of each biomarker (Supplemental Figures 3 - 12 and Supplemental Table 2) using the R package *sn*^*27*^.

### Modeling Application

#### Simulation Study

DPMs^8^ were trained on two randomly selected consecutive visits per participant (N=300), with the remaining six visits reserved for testing. We implemented a nonparametric bootstrap procedure (1,000 iterations). In each iteration, 80% of participants were sampled with replacement. Conversion from biomarker-negative to biomarker-positive was determined via linear interpolation between unmodeled visits when synthetic participants converted, yielding an observed time-from-positivity.

#### Observational Studies

We applied DPM to the available longitudinal biomarker data in a manner consistent with the simulation study. Supplemental Video 1 and 2 illustrate the modeling process in amyloid PET from DIAN and plasma pTau217/AB42 in ADNI. To ensure robust estimation of longitudinal change, we restricted analyses to individuals with baseline biomarker values above thresholds for reliable accumulation.^4^

### Model Performance Evaluation

#### Simulation Study

Model performance was quantified using mean absolute error (MAE) and root mean squared error (RMSE) between predicted and observed time-to-positivity in individuals who converted. Performance was evaluated across all considered ARC distributions and intra-individual noise levels, with selected representative exemplars presented in the primary text. Additional results are included as supplemental materials. Supplemental Videos 3 and 4 display the effects of increasing intra-individual noise on modeling. To quantify the contribution of distributional features to model error, we fit a three-way fixed-effects analysis of variance (ANOVA), including main effects (the central tendency, skew, and spread) and their interactions.

Model stability was assessed by examining the distribution of predicted biomarker timing MAEs across simulated intra-individual variability levels.

#### Observational Studies

We evaluated DPM performance using both MAE and RMSE. Because there were few observed conversion events, we also performed a supplemental model calibration analysis, where we evaluated the proportion of individuals who were always biomarker positive or biomarker negative based on their baseline predictions. We performed a survival analysis with age at first amyloid positive visit as the event, calculating the Harrell’s concordance index (C-index) to evaluate how well the order of amyloid time forecasts based on baseline visit aligned with the true order of age at conversion. Finally, to assess how informative biomarker levels are for predicting individual level disease timing, we examined bootstrapped distributions of Z scores at key timepoints (*t=-2,0,4,6,8*) relative to biomarker positivity.

## RESULTS

### Data Characterization

#### Simulation Study

We first defined inter-individual variability by perturbing the mathematical parameters that define a skew-normal distribution. We perturbed the central tendency (ξ; ∈ {−1, 0, 1}), spread (ω; ∈ {0.5, 1, 1.5, 2}), which increases the heterogeneity of ARC values, and skew (α; ∈ {−2, 0, 2}), which increases the number of individuals with high ARC values (Figure 2A, Supplemental Figure 2).

**Figure 2.**
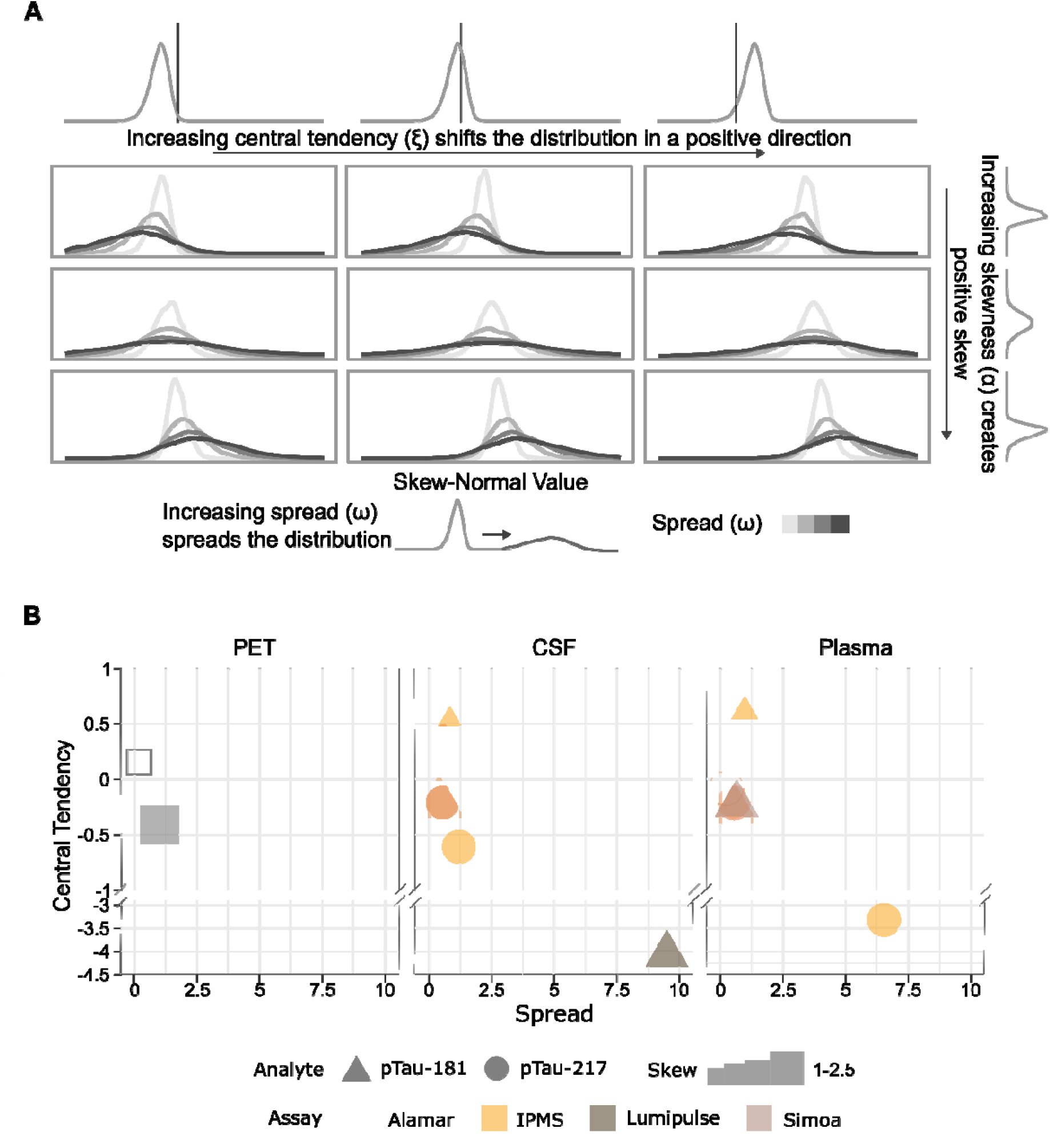
Rate of Change Distribution characteristics. **A.**Distribution visualization of the synthetic data generated. We performed a full grid combination of biologically plausible values for skew (α), central tendency (ξ), and spread (ω) **B**. Characterization of the annualized rates of change’s distribution for each of the longitudinal biomarkers measured/available in the DIAN (solid shapes) and ADNI (open shapes) studies.

#### Observational Studies

After defining the necessary thresholds for biomarker positivity and reliable accumulation, we characterized the three parameters that define a skew-normal distribution in the DIAN and ADNI datasets (Figure 2B; Supplemental Figures 3 – 17, Supplemental Tables 3 and 4). The underlying correlational structure between synthetic and observational data was consistent (Supplemental Figure 1). Most CSF-derived biomarkers had low skew, indicating a high number of individuals with negative or low-but-positive ARCs. In contrast, most plasma biomarkers showed higher skew, suggesting a greater abundance of individuals above the threshold for reliable accumulation; however, plasma biomarkers tended to lower central tendency values (ξ) which would increase the prevalence of negative ARC cases. This is undesirable because DPM assumes biomarker levels exclusively increase over time. Amyloid PET in DIAN had the highest skew value, consistent with the largest number of individuals with positive ARCs. Amyloid PET in ADNI had a skew value much more similar to the fluid biomarkers, but a somewhat higher central tendency potentially due to greater site heterogeneity across the study. In both cases, amyloid PET had positive correlations between baseline value and ARC (Supplemental Figure 1). Finally, most biomarkers showed small spread values, indicating low inter-individual variability. There are two exceptions to these parameters: plasma %p-tau217 measured using IP-MS, and CSF pTau181 measured using Lumipulse. Both showed relatively low central tendency with higher spread. This suggests that there are a limited number of individuals with very high ARC, which is a desirable characteristic for DPM, while simultaneously having many individuals below the threshold for reliable accumulation (not likely to have positive ARC), which is not desirable.

### DPM Evaluation

#### Group Level Performance

Our results suggest that the best group level DPM performance occurs when the central tendency is high, corresponding to a scenario where most individuals have higher ARCs; the spread is low, corresponding with minimal inter-individual variability/ less heterogeneity in accumulation rate; and the skew is high, which suggests a subset of individuals exist with exceptionally high rates of accumulation (Figure 3A, Supplemental Figure 1B).

**Figure 3.**
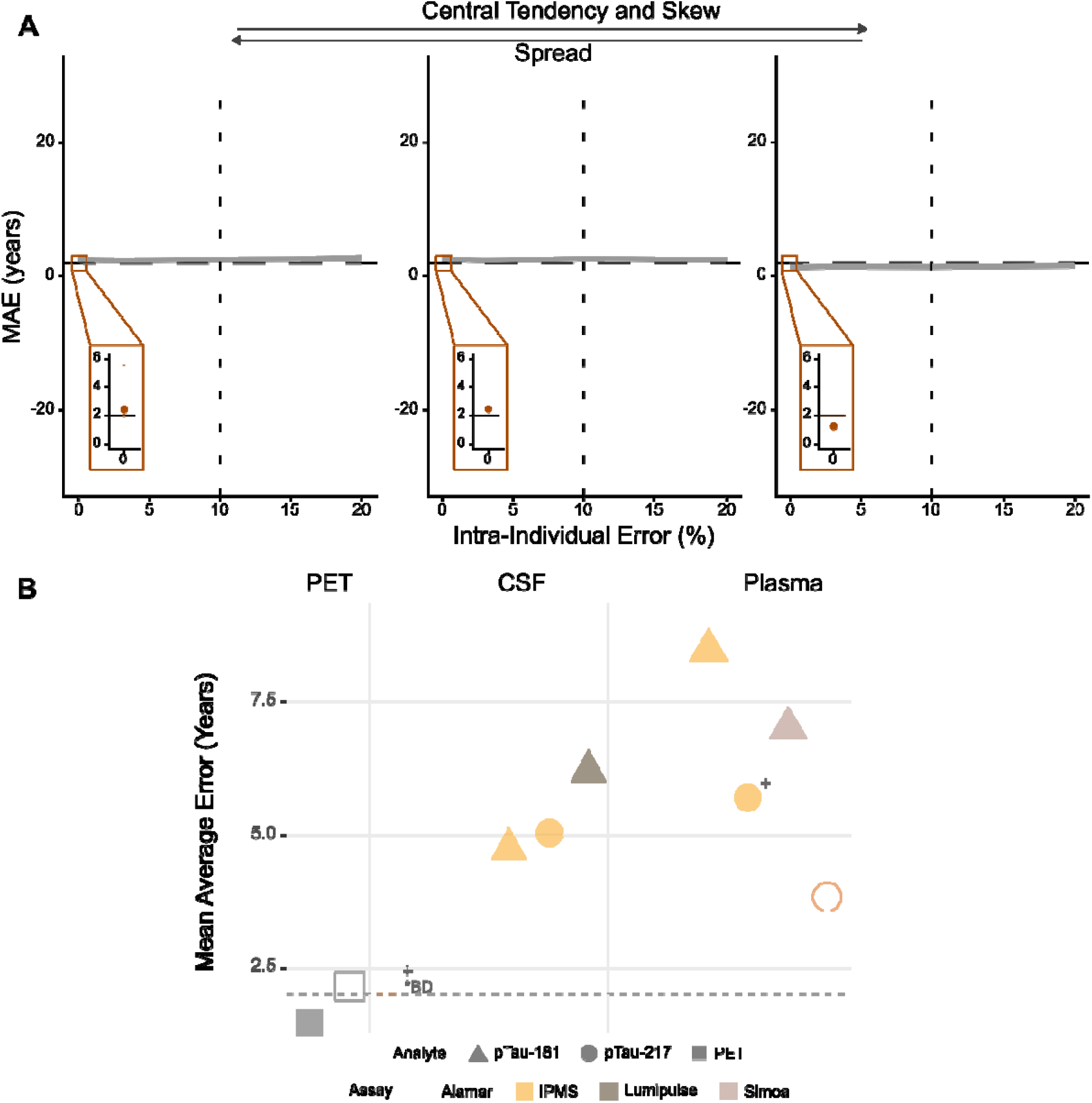
Group Level Performance Results. **A**. As intra-individual variability increased, model stability declined in synthetic datasets. Substantial widening of confidence intervals was observed when variability exceeded approximately 10–15%, indicating reduced model stability. Higher skew (α), lower spread (ω), and higher central tendency (ξ) were associated with lower MAE. Best-performing configurations approach the study sampling frequency (dashed line). Inset panels show a detailed view of the error performance in the case of 0% within-individual variability. **B**. Scatter plot depicting the Mean Average Error of the amyloid biomarkers available in the DIAN (solid shades) and ADNI (open shades) cohorts. Amyloid PET and Alamar CSF brain derived phosphorylated tau 217 (BD-p-tau217) demonstrated MAE below the study’s effective sampling frequency threshold (dashed line), indicating good performance in observed biomarker data. ^*t*^*Validation samples with documented conversion from biomarker-negative to biomarker-positive status were limited (n<5) and estimates of MAE should be interpreted with caution.*

The results of the fixed ANOVA highlighted that no single skew-normal distribution characteristic had a disproportionate influence on model performance, instead, the three-way interaction of these terms explained 45% of the variance, suggesting that DPM performance is affected by the ARC distribution characteristics wholistically, rather than specifically by one key parameter.

To examine the impact of intra-individual variability, we again generated distributions based on a range of skew-normal parameters and then added noise ranging from 3% to 20%, aligning with literature-derived estimates of biomarker test-retest variability^16,17,19^. When evaluating model performance, the MAE ranged from 1.30 to 2.64 years, regardless of skew-normal and intra-individual variability characteristics. This is near the sampling frequency of the synthetic data and suggests limited impact on model performance. However, the stability of the predictions, as highlighted by the 95% bootstrapped confidence intervals, were highly sensitive to intra-individual variability (Figure 3B).

We applied DPM (Supplemental Figures 18, 19) and calculated model performance in cases of observed conversion. PET demonstrated the highest accuracy across both DIAN (MAE=1.64 years; RMSE=2.34 years) and ADNI (MAE=2.15; RMSE=3.08) (Figure 3B). Among the fluid biomarkers, the best performing biomarkers in DIAN where at least five validations samples were available was IP-MS CSF %p-tau181 (MAE=4.80 years, RMSE=5.88 years) for CSF and plasma p-tau181 measured using Simoa (MAE=6.58 years, RMSE=8.14 years). We expected Alamar CSF BD p-tau217 and IP-MS plasma %p-tau217 to have a high degree of fidelity to amyloid PET. However, the number of conversion events was very limited, making it challenging to assess their accuracy. Further, the characteristics of the ARC for these two biomarkers are not different from the other fluid biomarkers, suggesting that quality of performance of these biomarkers, if distinct from other fluid biomarkers, would be due to lower intra-individual variability. Group-level performance in the available ADNI plasma p-tau217/Aβ42 was relatively stronger than performance in the DIAN plasma and CSF biomarkers (MAE=3.78, RMSE=4.88 in ADNI vs. MAEs of 4.2–9.1 and RMSEs of 5.2–10.8 years). This difference was not driven by differences in sample size (N=108 compared to N=36-475), and thus may suggest lower within-individual variation in the p-tau217/Aβ42 ratio. Unfortunately, due to the unavailability of test-retest data in the research cohorts, we were not able to confirm it. Model calibration evaluation demonstrated that DPM exhibits consistently good performance in individuals who have already converted to biomarker positive; however, in individuals who were negative at study enrollment, forecast quality is less certain (Supplemental Figure 20).

#### Individual Level Performance

For the synthetic data, the most distinct distributions of biomarker levels were associated with specific temporal estimates in the case of high central tendency and high spread values, with skew values having less impact (Figure 4A). The highlighted box in Figure 4A is suggestive of strong individual performance. This plot visualizes the distribution of Z score values that are associated with time predictions −2, 0, 2, 4, and 6 years based on the 1000 iteration bootstrap in the case of 0% intra-individual noise. The distinct distributions indicate that across iterations, there is a relatively specific association between a given biomarker level and its corresponding temporal prediction. In contrast, the distributions associated with high levels of intra-individual noise (righthand side of Figure 4A) are overlapping, suggesting that for a given biomarker value, depending on the sample that the model is fit on, may yield estimates of a wide range of temporal estimates with nearly equal probability. If the temporal estimate is highly dependent on the sample chosen for model fitting, the model is not robust and thus is unlikely to yield valid individual estimates

**Figure 4.**
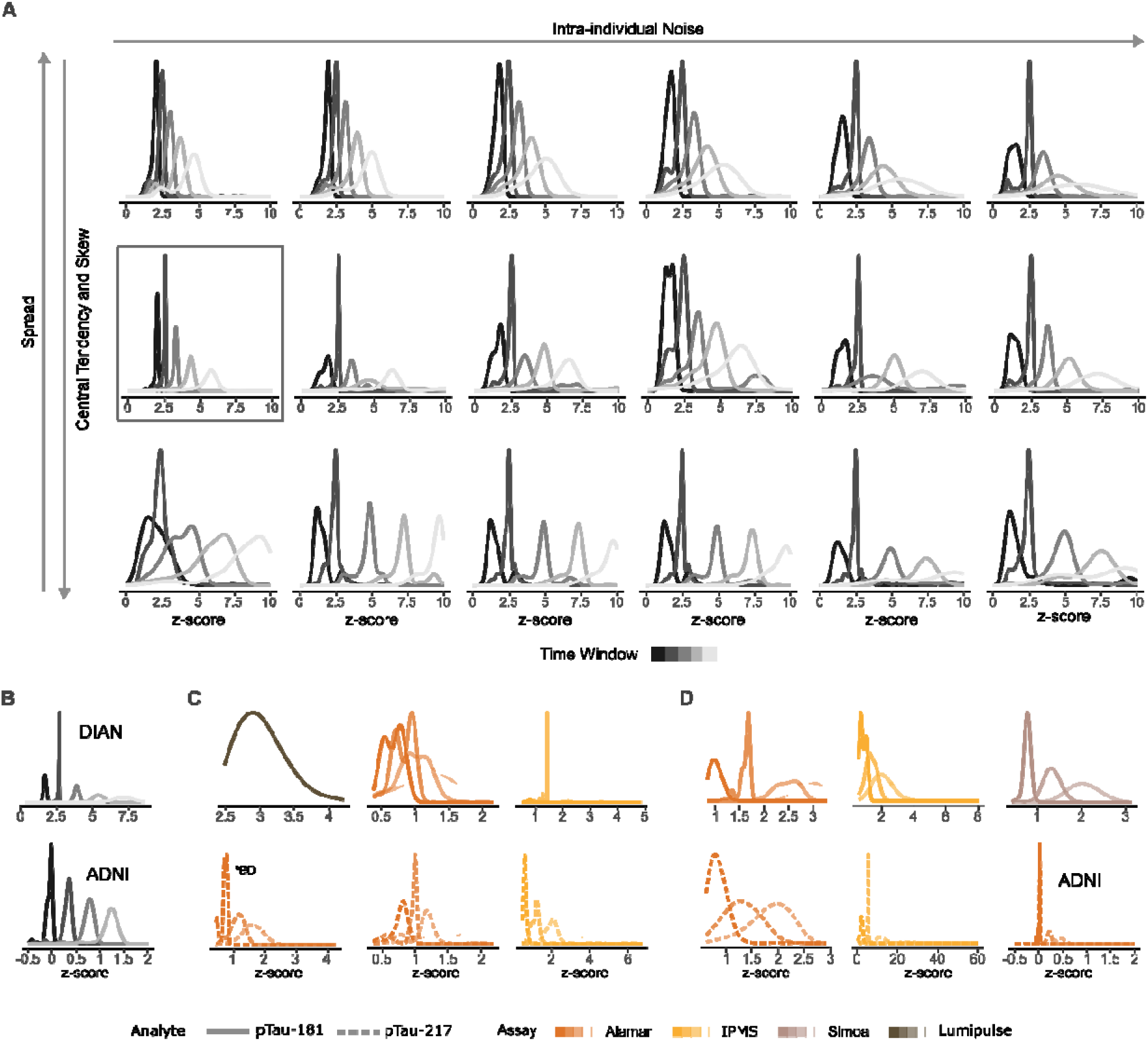
Performance at Individual Level Performance Assessment. When we consider model specificity, the distribution of biomarker levels that are specifically associated with a time window are more likely to perform well at the individual level. **A**. Summary of synthetic data results. Increasing intra-individual variability leads to decreased model specificity, whereas perturbing central tendency or skew has much less effect. The highlighted box indicates ideal performance, where distinct distributions can be seen at two year sampling intervals, suggesting that specific biomarker values have strong associations with specific temporal predictions. **B**. Amyloid PET performance has the strongest specificity with clearly separated ARC z-score distributions in both observational studies. **C**. CSF biomarkers demonstrate considerable overlap, regardless of assay or tau phosphorylation site. In the case of CSF pTau181 – Lumipulse, only one distribution is generated because the threshold for reliable accumulation is so great that DPM predictions do not begin until near time estimate of +6 years. **D**. Plasma biomarkers also generally exhibited characteristics of unstable estimates across iterations, where the distributions of biomarker levels associated with a specific temporal estimate overlapped considerably. For plasma pTau217/Aβ42, the distribution of Z scores at the time of conversion are tightly coupled, suggesting specificity at the time of conversion; however, at higher z score levels, the distributions are wide suggesting that models would not yield consistent temporal estimates.

Low spread values often resulted in poor individual performance. Low spread paired with low central tendency and low skew leads to a very high proportion of individuals below the meaningful threshold for ARC. These individuals may not be accumulating amyloid, or, if they are, they have not accumulated sufficient pathology to be above the measurement floor threshold of the biomarker of interest. These individuals cannot be modeled using DPM. As we increased intra-individual noise, distribution distinctness decreased, meaning that specific biomarker levels did not robustly map onto specific temporal estimates (Figure 4A).

When evaluating model performance at the individual level, only amyloid PET displayed a high degree of distributional specificity in both DIAN and ADNI (Figure 4B). This specificity suggests that for a given measurement, the DPM would predict the same temporal estimate (i.e. disease stage) consistently across bootstrapped iterations, mapping the biomarker level for a given individual onto a specific temporal estimate. Other models, such as Alamar CSF p-tau181, show a high degree of distribution overlap (Figure 4C). This is also true for CSF BD-ptau217 and plasma IP-MS %p-tau217, among others (Figure 4D). Notably, the application of the reliable accumulation heuristic makes it such that biomarkers such as Lumipulse CSF p-tau181 and Simoa plasma p-tau181 estimate time relative to conversion only years after conversion. The relative accumulation threshold associated with meaningful signal increase in the biomarker is a fundamental characteristic to consider when applying DPM. This finding is further supported by our calibration study, which found much worse DPM performance in individuals who were amyloid negative throughout the study than in individuals who were amyloid positive throughout (Supplemental Figure 20). The plasma p-tau217/Aβ42 values from ADNI yield relatively high specificity near conversion (times=0,2), but specificity declines at later stages, suggesting that elevated assay values do not reliably separate out into distinct temporal estimates.

The survival analysis revealed that for all biomarkers, good concordance (>0.80) was attained, suggesting that the relative ordering of participants was largely preserved by the DPM (Supplemental Figure 21). C-index results suggest that DPM has good utility for disease staging (defining individuals as early vs. late converters within research cohort) for all biomarkers, even if inspection of distributions suggests it does not provide consistently specific estimates of actual time from positivity across biomarkers.

## DISCUSSION

This study systematically evaluated how measurement variability and biological heterogeneity influence DPM. Using simulations informed by longitudinal research data, we found that these sources of variability differentially affect performance. Low within-individual variability improved the precision of individual temporal estimates, whereas favorable longitudinal accumulation characteristics reduced group-level estimation error. We validated these observations in autosomal dominant AD (DIAN) and sporadic AD (ADNI), identifying biomarker properties most conducive to temporal inference. Across all biomarkers examined, only amyloid PET estimated time from amyloid positivity with error approaching typical study sampling intervals.

A prerequisite for DPM is defining a biologically meaningful reference point for disease onset. Rather than relying solely on observed conversion events, which are concentrated near study enrollment, we used simulations spanning the full preclinical disease course to evaluate performance across a 20-year interval. In observational data, we defined time zero using the biomarker threshold that best discriminated individuals above and below 18 Centiloids^25^. Consistent with prior work, CSF biomarkers discriminated amyloid positivity better than plasma biomarkers ^28^, likely reflecting their closer relationship to central nervous system pathology ^29^. Reliable amyloid PET accumulation occurred below the positivity threshold (12.6 in DIAN, 13.2 CL in ADNI), indicating that meaningful signal exists before overt amyloid positivity. In contrast, several fluid biomarkers exhibited accumulation thresholds above their positivity thresholds, suggesting that reliable temporal inference begins only after amyloid conversion. While potentially problematic for modeling, this is consistent with the previously reported observation that tau phosphorylation sites reliably increase after 30 CL and that p-tau181 increases later than p-tau217^30^.

Longitudinal accumulation-rate distributions strongly influenced DPM performance. Biomarkers with lower error exhibited higher average accumulation, lower between-individual variability, and positively skewed distributions, resulting in larger proportions of individuals with consistently positive accumulation. Simulations suggested that DPM performs best when individuals either exhibit minimal accumulation or relatively homogeneous positive accumulation. Broad distributions weaken the relationship between biomarker value and disease timing because individuals with similar biomarker values may accumulate pathology at markedly different rates, while negative accumulation rates violate the fundamental assumption that biomarkers increase monotonically. These characteristics explain the superior performance of amyloid PET in both cohorts and plasma p-tau217/Aβ42 in ADNI. Cohort differences in PET accumulation likely reflect acquisition and harmonization differences, emphasizing that DPM performance depends not only on biomarker modality but also dataset-specific longitudinal characteristics.

At the individual level, intra-individual variability was the dominant determinant of model stability. Increasing measurement noise produced only modest increases in median error but substantially widened confidence intervals, and simulations showed that distributions corresponding to individual temporal estimates became poorly separated once noise exceeded approximately 10–15%. This pattern mirrors empirical observations: amyloid PET and CSF biomarkers, which have lower test–retest variability ^16,17^, showed greater temporal specificity than plasma biomarkers. This may also reflect biological differences. Amyloid PET measures cumulative pathological burden and is therefore highly autoregressive, which may improve DPM estimation. CSF biomarkers generally outperformed plasma biomarkers, which mayreflect their closer relationship to central nervous system pathology and reduced peripheral influence.

Plasma p-tau217/Aβ42 showed good performance at the group level but showed reduced specificity beyond the earliest periods following conversion. This pattern is consistent with saturation effects that constrain temporal inference at higher biomarker levels. Prior studies have similarly shown plateauing of CSF tau phosphorylation markers between 50–90 Centiloids^31,32^ and saturation of plasma p-tau181^13^ at later disease stages. Therefore, DPM is fundamentally bounded at the lower end by the relationship between accumulation and positivity thresholds and at the upper end by biomarker saturation.

These findings help explain why published amyloid-clock models have largely been interpreted at the group level^2,3,8^. Current appropriate-use recommendations for AD biomarkers limit clinical biomarker testing to individuals with objective cognitive impairment undergoing diagnostic evaluation, and do not recommend biomarker testing in cognitively unimpaired individuals outside of research studies or clinical trials. Correspondingly, clock models are not currently used in routine clinical care or for participant selection in trials, which instead rely on biomarker positivity thresholds. Our findings demonstrate that inter-individual heterogeneity, measurement variability, and biomarker dynamic range independently constrain individualized temporal inference, providing a mechanistic explanation for these current practices.

These findings may extend to biomarker interpretations outside the context of amyloid clocks. Because amyloid clocks are simply a transformation of both baseline value and longitudinal rate of change, this study offers key insights into longitudinal performance of biomarkers more broadly. Our findings support previous assertions that biomarkers with high test-retest variability may have limited utility for accurately locating an individual’s position on the pathological continuum^33^.

This study has several limitations. Data availability differed substantially across biomarker modalities, with amyloid PET having greater longitudinal availability than many fluid biomarkers. As a result, sample sizes varied considerably across analyses. To partially address this, we subsampled the ADNI dataset. Several biomarkers had limited numbers of observed conversion events, restricting performance evaluation. Presented MAE and RMSE values are likely underestimates, as data from research cohorts is biased to have conversion events occurring in close proximity to scan time. This was managed, to the best of our ability, by performing the supplemental model calibration evaluation in the research data and through simulation studies, which allowed us to have much larger temporal separation between conversion event and scan collection. Test–retest variability estimates were derived from the literature because sufficiently dense repeat-measurement data were not available within the cohorts themselves. Most evaluated biomarkers remain research-use only, and variability in fluid biomarkers may additionally reflect batch effects. Although ADNI included clinically relevant fluorine-18 tracers, DIAN relied on PiB PET, which is not used clinically and may exhibit different longitudinal properties than clinically deployed tracers. Finally, these analyses relied on a single disease-progression modeling framework, whereas multiple DPM approaches exist in the literature^3,6^ and may differ in performance characteristics.

In summary, DPM depends on three fundamental biomarker properties: dynamic range, inter-individual heterogeneity, and intra-individual variability. Floor and ceiling effects constrain inference at early and late disease stages. Accuracy depends on relatively homogeneous rates of change across individuals. Within individual noise impacts model stability, weakening the specific associations between biomarker level and temporal estimate and reinforcing that DPM is primarily a group-level tool. Future applications of DPM should carefully consider the distribution of individual accumulation rates and measurement reliability when selecting biomarkers for temporal modeling. We conclude that DPM is more robust for describing group-level trajectories than for estimating precise timelines in individuals and resampling strategies to quantify model stability and mitigate the influence of outliers should be considered in future applications of DPM.

## Supporting information

Supp Video - DIAN CSF pT181

Supp Video - DIAN PET PiB

Supp Video - 3% Within Individ Var

Supp Video - 20% Within Individ Var

## DATA AVAILABILITY STATEMENT

Consistent with best practices, code used to generate all simulations is available at github.com/jwisch/AmyloidClocks_SimandDIAN. For reproducibility, random seeding is documented there as well. DIAN data is available upon request via https://dian.wustl.edu/dian-observational-data-request-form/. ADNI data is available on LONI.

## References

1. Oxtoby, N. Data-Driven Disease Progression Modeling. in Machine Learning for Brain Disorders (ed. Colliot, O.) vol. 197 511–532 (Humana Press, Saskatoon, Canada, 2023).

2. Birdsill, A. C. et al. Trajectory of clinical symptoms in relation to amyloid chronicity. Alzheimer’s & Dementia: Diagnosis, Assessment & Disease Monitoring 14, (2022).

3. Betthauser, T. J. et al. Multi-method investigation of factors influencing amyloid onset and impairment in three cohorts. Brain 145, 4065–4079 (2022).

4. Wisch, J. K. et al. Comparison of Amyloid Chronicity and EYO in Autosomal Dominant Alzheimer Disease. Alzheimer’s & Dementia (2025).

5. Milà-Alomà, M. et al. Timing of Changes in Alzheimer’s Disease Plasma Biomarkers as Assessed by Amyloid and Tau <scp>PET</scp> Clocks. Ann. Neurol. 98, 508–523 (2025).

6. Petersen, K. K. et al. Predicting onset of symptomatic Alzheimerl1s disease with plasma p-tau217 clocks. Nat. Med. https://doi.org/10.1038/s41591-026-04206-y (2026) doi:10.1038/s41591-026-04206-y.

7. Jack, C. R. et al. Revised criteria for diagnosis and staging of Alzheimer’s disease: Alzheimer’s Association Workgroup. Alzheimer’s & Dementia https://doi.org/10.1002/alz.13859 (2024) doi:10.1002/alz.13859.

8. Budgeon, C. A. et al. Constructing longitudinal disease progression curves using sparse, short-term individual data with an application to Alzheimer’s disease. Stat. Med. 36, 2720–2734 (2017).

9. Schindler, S. E. et al. Predicting Symptom Onset in Sporadic Alzheimer Disease With Amyloid PET. Neurology 97, (2021).

10. De Meyer, S. et al. Plasma pTau181 and pTau217 predict asymptomatic amyloid accumulation equally well as amyloid PET. Brain Commun. 6, (2024).

11. Benedet, A. L. et al. The accuracy and robustness of plasma biomarker models for amyloid PET positivity. Alzheimers Res. Ther. 14, 26 (2022).

12. Salvadó, G. et al. Specific associations between plasma biomarkers and postmortem amyloid plaque and tau tangle loads. EMBO Mol. Med. https://doi.org/10.15252/emmm.202217123 (2023) doi:10.15252/emmm.202217123.

13. Moscoso, A. et al. Time course of phosphorylated-tau181 in blood across the Alzheimer’s disease spectrum. Brain 144, 325–339 (2021).

14. Schindler, S. E. et al. Head-to-head comparison of leading blood tests for Alzheimer’s disease pathology. Alzheimer’s & Dementia 20, 8074–8096 (2024).

15. Ashton, N. J. et al. The Alzheimer’s Association Global Biomarker Standardization Consortium (GBSC) plasma phospho-tau Round Robin study. Alzheimer’s & Dementia 21, (2025).

16. Tolboom, N. et al. Test-retest variability of quantitative [11C]PIB studies in Alzheimer’s disease. Eur. J. Nucl. Med. Mol. Imaging 36, 1629–1638 (2009).

17. Jonaitis, E. M. et al. CSF Biomarkers in Longitudinal Alzheimer Disease Cohorts: Pre-Analytic Challenges. Clin. Chem. 70, 538–550 (2024).

18. Brum, W. S. et al. Biological variation estimates of Alzheimer’s disease plasma biomarkers in healthy individuals. Alzheimer’s & Dementia 20, 1284–1297 (2024).

19. Cullen, N. C. et al. Test-retest variability of plasma biomarkers in Alzheimer’s disease and its effects on clinical prediction models. Alzheimers Dement. 19, 797–806 (2023).

20. Gong, K. et al. High-sensitivity plasma proteomics reveals disease-specific signatures and predictive biomarkers of Alzheimer’s disease phenotypes in a large mixed dementia cohort. Preprint at https://doi.org/10.21203/rs.3.rs-6440485/v1 (2025).

21. Arranz, J. et al. Diagnostic performance of plasma pTau217, pTau181, Aβ1-42 and Aβ1-40 in the LUMIPULSE automated platform for the detection of Alzheimer disease. Alzheimers Res. Ther. 16, 139 (2024).

22. Jagust, W. J. et al. The ADNI PET Core at 20. Alzheimer’s & Dementia 20, 7340–7349 (2024).

23. Shaw, L. M. et al. ADNI Biomarker Core: A review of progress since 2004 and future challenges. Alzheimer’s & Dementia 21, (2025).

24. Beer, J. C. et al. Longitudinal ComBat: A method for harmonizing longitudinal multi-scanner imaging data. Neuroimage 220, 117129 (2020).

25. Royse, S. K. et al. Validation of amyloid PET positivity thresholds in centiloids: a multisite PET study approach. Alzheimers Res. Ther. 13, 99 (2021).

26. Bollack, A. et al. Investigating reliable amyloid accumulation in Centiloids: Results from the AMYPAD Prognostic and Natural History Study. Alzheimer’s & Dementia 20, 3429–3441 (2024).

27. Azzalini, A. R package ‘sn’: The skew-normal and skew-t distributions (version 0. 4-18). Preprint at (2013).

28. Volluz, K. E. et al. Correspondence of CSF biomarkers measured by Lumipulse assays with amyloid PET. in 2021 Alzheimer’s Association International Conference (2021).

29. Roberts, K. F. et al. Amyloid-β efflux from the central nervous system into the plasma. Ann. Neurol. 76, 837–844 (2014).

30. Suárez-Calvet, M. et al. Novel tau biomarkers phosphorylated at T181, T217 or T231 rise in the initial stages of the preclinical Alzheimer’s continuum when only subtle changes in Aβ pathology are detected. EMBO Mol. Med. 12, (2020).

31. Wisch, J. K. et al. Predicting continuous amyloid PET values with CSF tau phosphorylation occupancies. Alzheimer’s & Dementia 20, 6365–6373 (2024).

32. Devanarayan, V. et al. Plasma pTau217 predicts continuous brain amyloid levels in preclinical and early Alzheimer’s disease. Alzheimer’s & Dementia https://doi.org/10.1002/alz.14073 (2024) doi:10.1002/alz.14073.

33. Ackley, S. F. et al. Substituting blood-based biomarkers for imaging measures in Alzheimer’s disease studies: implications for sample size and bias. J. Gerontol. A Biol. Sci. Med. Sci. 81, (2026).

