## Supplement for "Biomarker Variability Limits Individualized Amyloid Time Estimation in Alzheimer Disease"

**SUPPLEMENTAERY METHODS**

**Data Sources**

*Synthetic Data Generation*

Synthetic data were generated using custom R functions with fixed random seeds to ensure reproducibility (available at github.com/jwisch/AmyloidClocks_SimandDIAN). For each simulation, data were generated for 300 simulated individuals with randomly assigned baseline biomarker values. Baseline value selection was probabilistically weighted to enrich for participants who transitioned from below to above the biomarker positivity threshold within the follow-up period, thereby maximizing the number of conversion events available for model validation. Individual annualized rates of change (ARCs) were sampled from a skew-normal distribution parameterized by ξ (location), ω (scale), and α (skewness), rather than assuming normality, to better approximate empirical variability. Time intervals between measurements were sampled from a normal distribution consistent with observed observational intervals (mean = 2.2 years, sd = 1.06 years) in the DIAN data. Although missing data in the form of extended intervals between longitudinal sample collection is possible (e.g. a subject completed scanning in spring 2018, was scheduled to return in spring 2020 but was instead unable to complete scanning at that temporal interval and so instead completed scanning in spring 2022), this randomness associated with extended intervals between visits is represented in the synthetic data through the distribution sampling process. No “missing” values were interpolated in any dataset, synthetic or from observational research cohorts. For each simulated individual, longitudinal visits were generated for a total of eight visits.

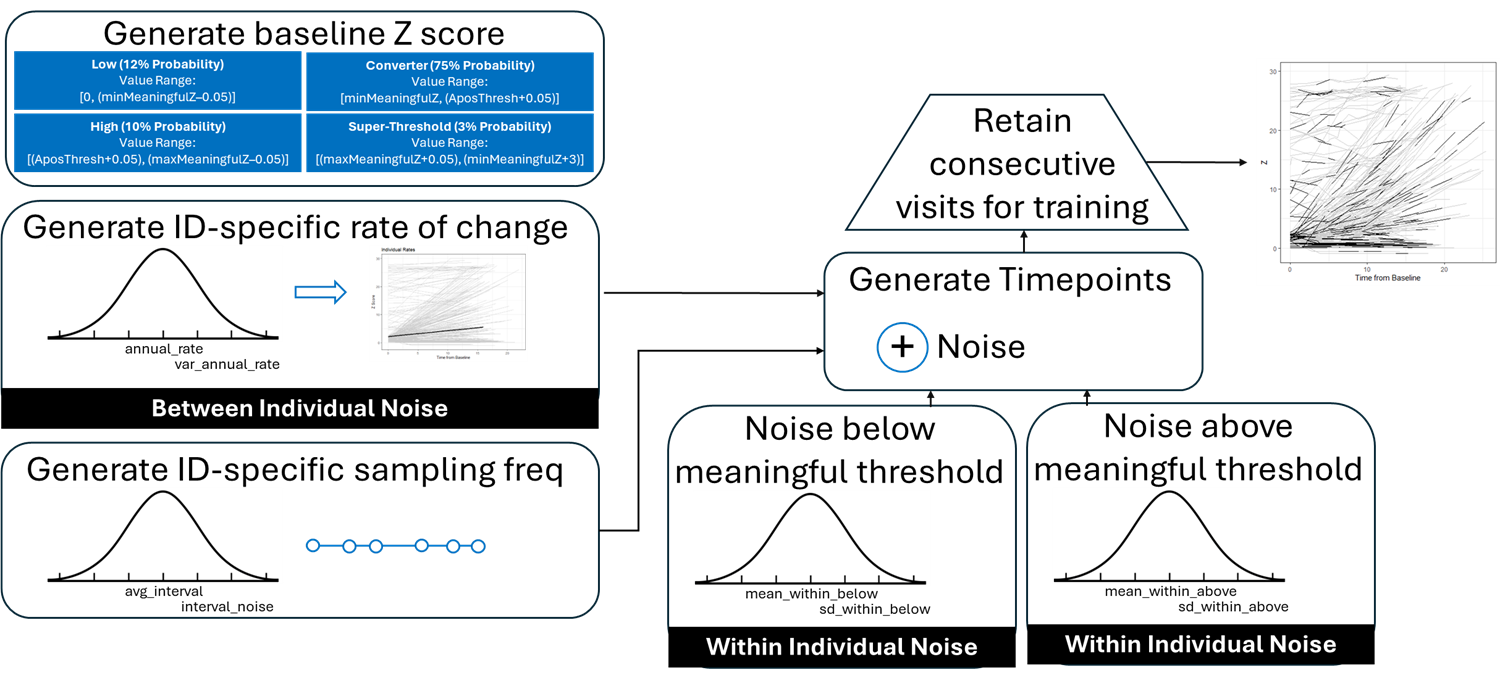

**Supp Methods Fig 1.** Method of synthetic data generation illustrated, where ID-specific annualized rates of change were drawn from a skew-normal distribution with characteristics defined by the skew normal parameters α, ξ, and ω and sampling frequencies were drawn from a normal distribution based on the characteristics of the DIAN sampling frequency. Within individual noise was injected between 0 and 15% as dictated by the simulation.

*Additional Details on Neuroimaging, DIAN*

Magnetic Resonance Imaging (MRI) was conducted using 3T Siemens or GE scanners. Reconstructed T1-weighted MR images were segmented (FreeSurfer 5.3, Desikan-Killiany atlas) and used as anatomical reference for PET. Dynamic amyloid-PET was obtained using ^11^C-Pittsburgh Compound B (PiB)^1,2^. Reconstructed PET time-series images were processed using the PET Unified Pipeline (PUP, https://github.com/ysu001/PUP)^1,3^. Briefly, images were smoothed to achieve a spatial resolution of 8 mm and corrected for inter-frame motion using in-house software^4–6^. The standardized uptake value ratio (SUVR) in each region was calculated with cerebellar grey as a reference region using the 40-70 minute post-injection window. A summary value for cortical amyloid^1^ was calculated as the arithmetic mean of the partial volume corrected (regional spread function^3,6^) SUVR of the precuneus and the superior frontal, rostral middle frontal, lateral orbitofrontal medial orbitofrontal, superior temporal and middle temporal regions^1^. SUVR summary values were transformed to Centiloids using previously published methods^7–9^. Individuals were considered amyloid positive if their cortical summary value exceeded 18 Centiloids (CL)^7^.

*Additional Details on Fluid Biomarkers, DIAN*

CSF and plasma biomarkers were collected at regular intervals from DIAN participants, as previously described^10^. Ten different phosphorylated tau (p-tau) isoforms measured in CSF and plasma were assayed. All measurements have been published (except plasma p-tau181measured using Simoa) and are available to qualified investigators via data request at: https://dian.wustl.edu/dian-observational-data-request-form/

Brefly, CSF p-tau181 was obtained using a Lumipulse G1200 Chemiluminescent Enzyme Immunoassay Assay platform as previously described^11^. Plasma and CSF p-tau181 and plasma p-tau217 were measured using immunoprecipitation-mass spectrometry (IP-MS) as previously described.^12^ ^13–15^ Additionally, plasma p-tau181 was measured with a Simoa assay developed at the University of Tuebingen (unpublished results).

The NULISAseq™ CNS Disease Panel 120 v1 was used to quantify CSF and plasma p-tau181, and p-tau217, following previously described protocols.^16–19^ The NULISAseq™ CNS Disease Panel 120 v1 reports values as NPQ units (NULISA Protein Quantification). These units are already standardized, normalized and log_2_ transformed. Samples that were below the detectability threshold were removed. NULISA quantification was produced in two batches and was thus scaled by plate to account for technical differences between the batches. To deal with batch effects in NULISA processing we performed principal component analyses (PCA) and further removed any outliers defined as any samples that were more than three standard deviations removed from the means of PC1 or PC2.

**Modeling Application**

*Inter-Individual Variability Simulation*

To evaluate the influence of different distributional assumptions, we performed a full grid search over skew-normal parameters: ξ ∈ {−1, 0, 1}, ω ∈ {0.5, 1, 1.5, 2}, and α ∈ {−2, 0, 2}, using the “*sn”* package in R ^20,21^ (Supplemental Figure 1A). The ranges were selected consistently with observed skew-normal parameters in the real world biomarker data. Synthetic datasets were then generated to simulate longitudinal biomarker trajectories under controlled conditions. Each dataset included 300 simulated individuals with eight visits, an average inter-visit interval of 2.02 ± 1.06 years, and ARC distribution parameters of mean 0.601 Z/year, SD = 0.934 Z/year) that we derived from the real world amyloid PET data. The minimum threshold for reliable accumulation was set at 0.902 Z (approximately 5 CL, based on the baseline measures of the full cohort), and the biomarker positivity threshold at 2.60 Z (18 CL).

*Intra-Individual Variability Simulation*

Intra-individual variability was modeled as a percentage of the observed value, with superthreshold noise levels varying between 3% and 20% (SD=3% of the mean), and subthreshold noise set to half that magnitude. These intra-individual variability levels were chosen to be consistent with characteristics of the real world data. PET test-retest variability has been reported to be as high as 8%^22^, CSF intra-individual noise may be between 5 and 10%^23^, and plasma-based estimates are closer to 20%^24,25^. We simulated datasets using a subset of the values from the inter-individuals noise simulation generated in the previous section (ξ ∈ {0, 1}, ω ∈ {0.5}, and α ∈ {0, 2}).

*Disease Progression Modeling Simulation*

DPMs^26^ were trained on two randomly selected consecutive visits per participant (N = 300), with the remaining six visits reserved for testing. Conversion from biomarker-negative to biomarker-positive was determined via linear interpolation between unmodeled visits, yielding a simulated true “time-to-positivity.” DPM performance was quantified by calculating mean absolute error (MAE) and root mean squared error (RMSE) between DPM-estimated time-to-positivity against simulated true values. All estimates were bootstrapped over 1,000 iterations, each using different visit pairs, and reported as median values with 95% confidence intervals.

To quantify the relative contribution of each parameter to model performance, we conducted a fully crossed factorial analysis across all combinations of ξ (location), ω (scale), and α (shape). After averaging MAE within each parameter combination to remove bootstrap variability, we fit a three-way fixed-effects ANOVA and decomposed the total sum of squares to estimate the proportion of systematic variance attributable to main effects and interactions.

To evaluate the specificity of a biomarker level for a given temporal estimate, which is an important consideration for individual-level application, we visualized the bootstrapped distribution of Z scores associated with temporal estimates of two years prior to conversion to biomarker positivity, at the conversion time, and two, four, and six years following conversion to biomarker positivity. We then performed a Bayesian estimation of the probability of temporal classification based on the observed distributions.

*Evaluation of Cohort-Derived Data (DIAN & ADNI)*

Cohort characteristics were summarized using medians and interquartile ranges for all demographic and clinical variables (Supplemental Table 1. ^~~15~~^Biomarker values were standardized to Z-scores relative to baseline distributions of non–mutation carriers in the DIAN cohort, providing a harmonized reference across modalities.

Biomarker positivity thresholds were defined separately for each biomarker using a bootstrapped cut-point optimization procedure^27^. Each of these biomarkers have historically been considered proxies for amyloid PET.^28–35^ Thresholds were derived by maximizing the Youden Index under the assumption of a ground truth biomarker positivity defined by 18 Centiloids (CL) on amyloid PET^7,36^. Model performance was evaluated using the area under the receiver operating characteristic curve (AUC), precision, and recall.

For each biomarker, ARCs were estimated using ordinary least squares regression. We performed two-component Gaussian Mixture Modeling on the distribution, defining “reliable accumulators” as individuals with a rate of change in the upper distribution^37^. This identified individuals with higher-than-typical rates of change, suggestive of a high degree of pathological change. We parameterized the empirical ARC distribution using a skew-normal model, yielding estimates of ξ (location), ω (scale), and α (shape) using the *sn* package in R^20,21^. To identify the minimum biomarker level consistent with reliable pathological accumulation^37^, we applied a bootstrapped cut-point analysis relating baseline biomarker values to ARCs, optimizing the Youden Index and reporting AUC, precision, and recall^27^.

*Application of DPM to Cohort-Derived Data (DIAN & ADNI)*

We applied DPM to the available biomarker data in DIAN and ADNI, similar to previous application^26^. However, for all longitudinal biomarker data, we only included individuals with a baseline biomarker value above the threshold for reliable accumulation.^38^ We implemented a nonparametric bootstrap procedure (1,000 iterations). In each iteration, 80% of participants were sampled with replacement, and individual biomarker trajectories were modeled using third-degree polynomial regression. We estimated the point at which each trajectory crossed the biomarker-positivity threshold, aligning trajectories by this crossing. Predicted biomarker values were interpolated across a standardized time window, and the median and 95% confidence intervals were computed, yielding a sample-level trajectory centered on the estimated time from biomarker-positivity.

To define the actual time from positivity, we used linear interpolation to estimate the actual age of conversion from negative to positive. Then we used the DPM model to forecast the age at conversion based on baseline biomarker value. ^38^ We calculated both MAE and RMSE between the model-predicted and interpolated conversion ages, consistent with the error metrics considered in the simulation study. We repeated the visualization of biomarker values associated with -2, 0, 2, 4, and 6 years relative to conversion to biomarker positivity to evaluate DPM specificity.

**SUPPLEMENTARY RESULTS**

**Data Characterization**

*Inter-Individual Variability*

We perturbed the location (ξ), scale (ω), and shape (α) parameters of the skew-normal distribution defining between-individual ARC variability. Increasing ξ shifts the distribution of ARC values in a positive direction, meaning that the central tendency for all individuals is to have a greater ARC. Increasing ω causes the distribution to spread, meaning there is greater heterogeneity in the ARC. Increasing α results in a positive skew, meaning that a larger number of individuals have relatively high ARC (Figure 2A, Supplemental Figure 1A).

Simulations revealed that higher location, lower scale, and higher shape were associated with lower model error at the group level (Figures 2A, Supplemental Figure 1B). Low location values impact the center of the distribution, causing the ARC distribution to fall below the biomarker threshold associated with meaningful accumulation. Low shape values results in a distribution that is left-skewed, which also correspond with an ARC distribution where most samples occur below the baseline biomarker threshold associated with reliable accumulation. Low scale values lead to a narrow distribution with low inter-individual variability. Low scale paired with low shape and low location leads to a very high proportion of individuals below the meaningful threshold for ARC, which is incompatible with successful DPM.

Consistent with the observation that higher location leads to better model performance, higher location was also associated with more specific “peaks” in a comparison of the distribution of Z_ARC_ values associated with disease time estimates of -2, zero (key event – biomarker conversion), two, four, and six years. Taller peaks and narrower distributions of Z_ARC_ values suggest that a given biomarker level is more specifically and uniquely associated with a disease time estimate. This specificity is necessary for extension to both individual-level modeling and cross-sectional data.

When we performed the post hoc ANOVA, we observed that fully 45% of the model variance across the full parameter grid search was explained by the three-way interaction among location, scale, and shape. In contrast, the sum of the three main effects was only 15% (location=6.6%, scale=3.0%, and shape=5.2%). Thus, model performance is driven by the combined associations between the three evaluated characteristics of the skew-normal distribution rather than by any parameter independently.

*Intra-Individual Variability*

Across presented simulation conditions, the median MAE ranged from 1.30 to 2.64 years when we compared the estimated time from amyloid positivity to the observed conversion to amyloid positivity, comparable to the average sampling interval of 2.02 years, indicating that intra-individual variability has limited impact on point estimates of time from amyloid positivity in DPM (Figure 3). However, model uncertainty was highly sensitive to increasing intra-individual variability, as evidenced by 95% confidence interval width. This effect was amplified in scenarios with greater inter-individual heterogeneity and right-skewed ARC distributions, demonstrating a meaningful interaction between intra- and inter-individual variability. Critically, once intra-individual variability reached typically10–15%, model estimates became generally unstable.

This model instability is reflected in the decreasing specificity of the distributions of Z-scored biomarker levels for a given time point (Figure 4). At low intra-individual variability, high peaks associated with relatively little variability in the distribution of Z values. This reflects the ideal behavior, where a given Z value is overwhelmingly associated with a single time value. At high intra-individual variability in the identified example, low peaks are associated with high variability in the distribution of Z values. This reflects low model specificity and is not ideal behavior for DPM, particularly for application at the individual level.

**Supplementary Methods References**

1.	Su, Y. *et al.* Quantitative analysis of PiB-PET with FreeSurfer ROIs. *PLoS One* **8**, (2013).

2.	Driscoll, I. *et al.* Correspondence between in vivo 11C-PiB-PET amyloid imaging and postmortem, region-matched assessment of plaques. *Acta Neuropathol.* **124**, 823–831 (2012).

3.	Su, Y. *et al.* Partial volume correction in quantitative amyloid imaging. *Neuroimage* **107**, 55–64 (2015).

4.	Hajnal, J. V. *et al.* A registration and interpolation procedure for subvoxel matching of serially acquired mr images. *J. Comput. Assist. Tomogr.* **19**, 289–296 (1995).

5.	Eisenstein, S. A. *et al.* Characterization of extrastriatal D2 in vivo specific binding of [ 18 F](N-methyl)benperidol using PET. *Synapse* **66**, 770–780 (2012).

6.	Rousset, O. G., Ma, Y. & Evans, A. C. Correction for partial volume effects in PET: Principle and validation. *Journal of Nuclear Medicine* **39**, 904–911 (1998).

7.	Royse, S. K. *et al.* Validation of amyloid PET positivity thresholds in centiloids: a multisite PET study approach. *Alzheimers Res. Ther.* **13**, 99 (2021).

8.	Collij, L. E. *et al.* Centiloid recommendations for clinical context‐of‐use from the AMYPAD consortium. *Alzheimer’s & Dementia* **20**, 9037–9048 (2024).

9.	Klunk, W. E. *et al.* The Centiloid project: Standardizing quantitative amyloid plaque estimation by PET. *Alzheimer’s and Dementia* **11**, 1-15.e4 (2015).

10.	Bateman, R. J. *et al.* Clinical and biomarker changes in dominantly inherited Alzheimer’s disease. *New England Journal of Medicine* **367**, 795–804 (2012).

11.	Llibre-Guerra, J. J. *et al.* Association of Longitudinal Changes in Cerebrospinal Fluid Total Tau and Phosphorylated Tau 181 and Brain Atrophy With Disease Progression in Patients With Alzheimer Disease. *JAMA Netw. Open* **2**, e1917126 (2019).

12.	Barthélemy, N. R., Horie, K., Sato, C. & Bateman, R. J. Blood plasma phosphorylated-tau isoforms track CNS change in Alzheimer’s disease. *Journal of Experimental Medicine* **217**, (2020).

13.	Barthélemy, N. R. *et al.* Site-Specific Cerebrospinal Fluid Tau Hyperphosphorylation in Response to Alzheimer’s Disease Brain Pathology: Not All Tau Phospho-Sites are Hyperphosphorylated. *Journal of Alzheimer’s Disease* **85**, 415–429 (2022).

14.	Barthelemy, N. *et al.* CSF tau phosphorylation occupancies at T217 and T205 represent improved biomarkers of amyloid and tau pathology in Alzheimer’s disease. *Nat. Aging* (2023).

15.	Barthélemy, N. R. *et al.* A soluble phosphorylated tau signature links tau, amyloid and the evolution of stages of dominantly inherited Alzheimer’s disease. *Nat. Med.* **26**, 398–407 (2020).

16.	Gong, K. *et al.* High-sensitivity plasma proteomics reveals disease-specific signatures and predictive biomarkers of Alzheimer’s disease phenotypes in a large mixed dementia cohort. Preprint at https://doi.org/10.21203/rs.3.rs-6440485/v1 (2025).

17.	Ibanez, L. *et al.* Benchmarking of a multi‐biomarker low‐volume panel for Alzheimer’s disease and related dementia research. *Alzheimer’s & Dementia* **21**, (2025).

18.	Ali, M. *et al.* Multi-cohort cerebrospinal fluid proteomics identifies robust molecular signatures across the Alzheimer disease continuum. *Neuron* **113**, 1363-1379.e9 (2025).

19.	Lin, W. *et al.* Abnormalities in core AD biomarkers precede inflammatory and glial markers in CSF in Autosomal Dominant Alzheimer’s Disease. Preprint at https://doi.org/10.64898/2026.03.31.26349851 (2026).

20.	Hossain, A. & Beyene, J. Application of skew-normal distribution for detecting differential expression to microRNA data. *J. Appl. Stat.* **42**, 477–491 (2015).

21.	Azzalini, A. R package ‘sn’: The skew-normal and skew-t distributions (version 0.4-18). Preprint at (2013).

22.	Tolboom, N. *et al.* Test-retest variability of quantitative [11C]PIB studies in Alzheimer’s disease. *Eur. J. Nucl. Med. Mol. Imaging* **36**, 1629–1638 (2009).

23.	Jonaitis, E. M. *et al.* CSF Biomarkers in Longitudinal Alzheimer Disease Cohorts: Pre-Analytic Challenges. *Clin. Chem.* **70**, 538–550 (2024).

24.	Brum, W. S. *et al.* Biological variation estimates of Alzheimer’s disease plasma biomarkers in healthy individuals. *Alzheimer’s & Dementia* **20**, 1284–1297 (2024).

25.	Cullen, N. C. *et al.* Test-retest variability of plasma biomarkers in Alzheimer’s disease and its effects on clinical prediction models. *Alzheimers Dement.* **19**, 797–806 (2023).

26.	Budgeon, C. A. *et al.* Constructing longitudinal disease progression curves using sparse, short‐term individual data with an application to Alzheimer’s disease. *Stat. Med.* **36**, 2720–2734 (2017).

27.	Thiele, C. *Package ‘Cutpointr’: Determine and Evaluate Optimal Cutpoints in Binary Classification Tasks*. https://cran.r-project.org/web/packages/cutpointr/cutpointr.pdf (2019).

28.	Zhong, X. *et al.* Plasma p‐tau217 and p‐tau217/Aβ1‐42 are effective biomarkers for identifying CSF‐ and PET imaging‐diagnosed Alzheimer’s disease: Insights for research and clinical practice. *Alzheimer’s & Dementia* **21**, (2025).

29.	Reimand, J. *et al.* Amyloid‐β PET and CSF in an autopsy‐confirmed cohort. *Ann. Clin. Transl. Neurol.* **7**, 2150–2160 (2020).

30.	Volluz, K. E. *et al.* Correspondence of CSF biomarkers measured by Lumipulse assays with amyloid PET. in *2021 Alzheimer’s Association International Conference* (2021).

31.	Alcolea, D. *et al.* Agreement of amyloid PET and CSF biomarkers for Alzheimer’s disease on Lumipulse. *Ann. Clin. Transl. Neurol.* **6**, 1815–1824 (2019).

32.	Palmqvist, S. *et al.* *Detailed Comparison of Amyloid PET and CSF Biomarkers for Identifying Early Alzheimer Disease*. (2015).

33.	Olsson, B. *et al.* CSF and blood biomarkers for the diagnosis of Alzheimer’s disease: a systematic review and meta-analysis. *Lancet Neurol.* https://doi.org/10.1016/S1474-4422(16)00070-3 (2016) doi:10.1016/S1474-4422(16)00070-3.

34.	Benedet, A. L. *et al.* The accuracy and robustness of plasma biomarker models for amyloid PET positivity. *Alzheimers Res. Ther.* **14**, 26 (2022).

35.	De Meyer, S. *et al.* Plasma pTau181 and pTau217 predict asymptomatic amyloid accumulation equally well as amyloid PET. *Brain Commun.* **6**, (2024).

36.	Molina‐Henry, D. P. *et al.* Racial and ethnic differences in plasma p‐tau217 ratio biomarker eligibility rates in a preclinical AD trial with lecanemab. *Alzheimer’s & Dementia: Diagnosis, Assessment & Disease Monitoring* **17**, (2025).

37.	Bollack, A. *et al.* Investigating reliable amyloid accumulation in Centiloids: Results from the AMYPAD Prognostic and Natural History Study. *Alzheimer’s & Dementia* **20**, 3429–3441 (2024).

38.	Wisch, J. K. *et al.* Comparison of Amyloid Chronicity and EYO in Autosomal Dominant Alzheimer Disease. *Alzheimer’s & Dementia* (2025).

**Supplemental Materials**

**Supplemental Table 1. Participant Demographics, DIAN**

|  | **Non-Mutation Carrier** | **Mutation Carrier** | p |
| --- | --- | --- | --- |
| Sample Size | 212 | 347 |  |
| Enrollment Age (mean (SD)) | 37.87 (11.39) | 38.24 (10.76) | 0.701 |
| Estimated Years to Symptom Onset (mean (SD)) | -11.28 (11.69) | -8.59 (11.17) | 0.007 |
| Sex (% Female) | 126 (59.4) | 191 (55.0) | 0.353 |
| Race (%) |  |  | 0.413 |
| American Indian or Alaska Native | 1 (0.5) | 0 (0.0) |  |
| Asian | 7 (3.3) | 18 (5.2) |  |
| Black or African American | 5 (2.4) | 4 (1.2) |  |
| Native Hawaiian or Other Pacific Islander | 0 (0.0) | 3 (0.9) |  |
| Other | 9 (4.2) | 14 (4.0) |  |
| Unknown | 3 (1.4) | 3 (0.9) |  |
| White | 187 (88.2) | 305 (87.9) |  |
| APOE Haplotype (%) |  |  | 0.54 |
| 22 | 3 (1.4) | 1 (0.3) |  |
| 23 | 23 (10.8) | 35 (10.1) |  |
| 24 | 7 (3.3) | 9 (2.6) |  |
| 33 | 118 (55.7) | 206 (59.4) |  |
| 34 | 58 (27.4) | 87 (25.1) |  |
| 44 | 3 (1.4) | 9 (2.6) |  |
| Family Mutation (%) |  |  | 0.257 |
| PSEN1 | 147 (69.3) | 261 (75.2) |  |
| PSEN2 | 22 (10.4) | 25 (7.2) |  |
| APP | 43 (20.3) | 61 (17.6) |  |
| Longitudinal Amyloid PET - PiB | 133 (62.7) | 210 (60.5) | 0.665 |
| Longitudinal CSF pT181 - IPMS | 179 (84.4) | 296 (85.3) | 0.875 |
| Longitudinal CSF pT181 - Lumipulse | 132 (62.3) | 218 (62.8) | 0.966 |
| Longitudinal Plasma pTau217 - IPMS | 29 (13.7) | 48 (13.8) | 1 |
| Longitudinal Plasma pTau181 - Simoa | 137 (64.6) | 215 (62.0) | 0.588 |
| Longitudinal CSF pT181 - IPMS | 102 (48.1) | 165 (47.6) | 0.967 |
| Longitudinal CSF pT217 - IPMS | 102 (48.1) | 165 (47.6) | 0.967 |
| Longitudinal Plasma pT181 - NULISA | 1 (0.5) | 83 (23.9) | <0.001 |
| Longitudinal Plasma pT217 - NULISA | 1 (0.5) | 83 (23.9) | <0.001 |
| Longitudinal CSF pT217 - NULISA | 0 (0.0) | 36 (10.4) | <0.001 |
| Longitudinal CSF pT181 - NULISA | 0 (0.0) | 36 (10.4) | <0.001 |
| Longitudinal CSF Brain Derived pT217 - NULISA | 0 (0.0) | 76 (21.9) | <0.001 |

**Supplemental Table 2. Participant Demographics, ADNI**

|  | **ADNI Participants** |
| --- | --- |
| N | 108 |
| Age at Baseline | 71.5 (7.3) |
| Gender (% Female) | 55 (50.9%) |
| Baseline Cortical Amyloid (CL) | 25.4 (37.8) |
| Baseline Plasma pT217 / AB42 | 0.01 (0.01) |
| Amyloid Positive by PET (% Positive) | 43 (39.8%) |
| Clinical Dementia Rating |  |
| 0 … | 67 (62.0%) |
| 0.5…. | 34 (31.5%) |
| 1…. | 7 (6.5%) |
| **All participants have longitudinal amyloid PET and plasma pT217/AB42 - Fujirubio* | |

**Supplemental Table 3.** Comparison of relevant biomarker characteristics for disease progression modeling, DIAN

| **Biomarker** | **N** | **Baseline visit threshold for Reliable Accumulation [Z]** | **Threshold for Amyloid Positivity [Z]** | **Mean ARC [Z/year]** | **Standard Deviation ARC** | **Central Tendency (ξ)** | **Spread**  **(ω)** | **Skew**  **(α)** |
| --- | --- | --- | --- | --- | --- | --- | --- | --- |
| Amyloid PET | 338 | 0.732 | 1.82 | 0.295 | 0.714 | -0.409 | 1.000 | 2.870 |
| CSF pTau217 [IP-MS] | 267 | 0.767 | 1.82 | 0.166 | 0.932 | -0.614 | 1.210 | 1.480 |
| CSF pTau217 [NULISA] | 237 | 0.469 | 0.27 | 0.084 | 0.341 | 0.398 | 0.046 | -1.760 |
| Plasma pTau217 [NULISA] | 84 | 0.332 | 0.51 | 0.111 | 0.396 | -0.191 | 0.496 | 1.220 |
| Plasma pTau217 [IP-MS] | 77 | 3.294 | 3.87 | 0.802 | 5.097 | -3.320 | 6.530 | 1.390 |
| CSF pTau181 [IP-MS] | 264 | 2.882 | 1.37 | -0.035 | 0.616 | 0.529 | 0.834 | -1.770 |
| CSF pTau181 [NULISA] | 237 | 0.297 | 0.46 | 0.0399 | 0.286 | 0.288 | 0.378 | -1.540 |
| Plasma pTau181 [NULISA] | 84 | -2.018 | -1.8 | 0.135 | 0.479 | -0.231 | 0.600 | 1.220 |
| CSF pTau181 [Lumipulse] | 348 | 0.699 | 1.12 | 1.02 | 8.031 | -4.070 | 9.500 | 0.924 |
| Plasma pTau181 [IP-MS] | 77 | 1.683 | 1.38 | 0.003 | 0.76 | 0.609 | 0.968 | -1.350 |
| Plasma pTau181 [Simoa] | 350 | 1.51 | 0.36 | 0.126 | 0.547 | -0.225 | 0.649 | 0.941 |

**Supplemental Table 4.** Comparison of relevant biomarker characteristics for disease progression modeling, ADNI

| **Biomarker** | **N** | **Baseline visit threshold for Reliable Accumulation [Z]** | **Threshold for Amyloid Positivity [Z]** | **Mean ARC [Z/year]** | **Standard Deviation ARC** | **Central Tendency (ξ)** | **Spread**  **(ω)** | **Skew**  **(α)** |
| --- | --- | --- | --- | --- | --- | --- | --- | --- |
| Amyloid PET | 108 | -0.151 | 0.013 | 0.065 | 0.17 | 0.155 | 0.193 | -0.728 |
| Plasma pTau217 / AB42 [Fujirebio] | 108 | -0.018 | 0.026 | 0.072 | 0.19 | -0.0833 | 0.242 | 1.600 |

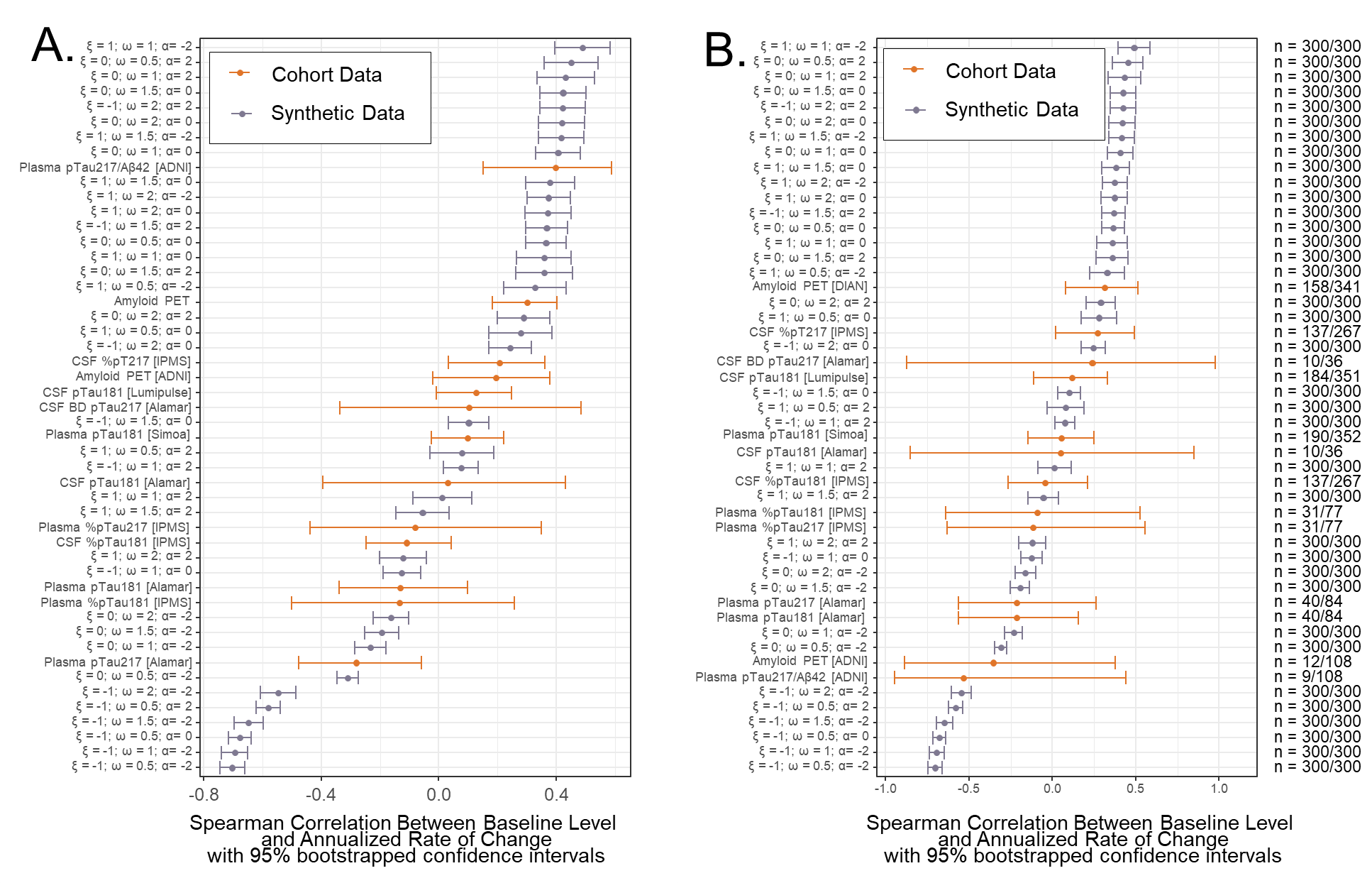

**Supplemental Figure 1.** The generated synthetic data has varying degrees of correlation between baseline level and annualized rate of change (ARC). We performed 500 iteration bootstraps of Spearman Correlations to estimate the correlation between baseline biomarker level and ARC of all considered synthetic datasets and the research cohort data. The underlying structure of the associations between baseline biomarker and ARC are consistent except in the cases of very low central tendency in the synthetic datasets (ξ = -1), which yields highly negative correlations between baseline and ARC. This negative association is inconsistent with DPM assumptions. Of note, the best performing datasets obtained from research cohorts (Amyloid PET, DIAN & ADNI; plasma pTau217/AB42, ADNI; CSF pT181, DIAN) have the highest positive correlations between baseline value and ARC, pointing to another heuristic that could be deployed to identify biomarkers likely to yield good results under DPM assumptions (A). Because correlations may be impacted by the influence of differences in sampling frequency on the calculation of ARC, we repeated the analysis limiting the ARC calculation to only include 2 samples per individual, and requiring that those samples be collected between 1.8 and 2.5 years apart. The results were largely consistent, with the exception of the ADNI data. These correlations were lower in the subsetted analysis because the majority of samples were collected farther apart. In the righthand column we show the number of samples used in the subsetted calculation relative to the total number of samples used in the original calculation.

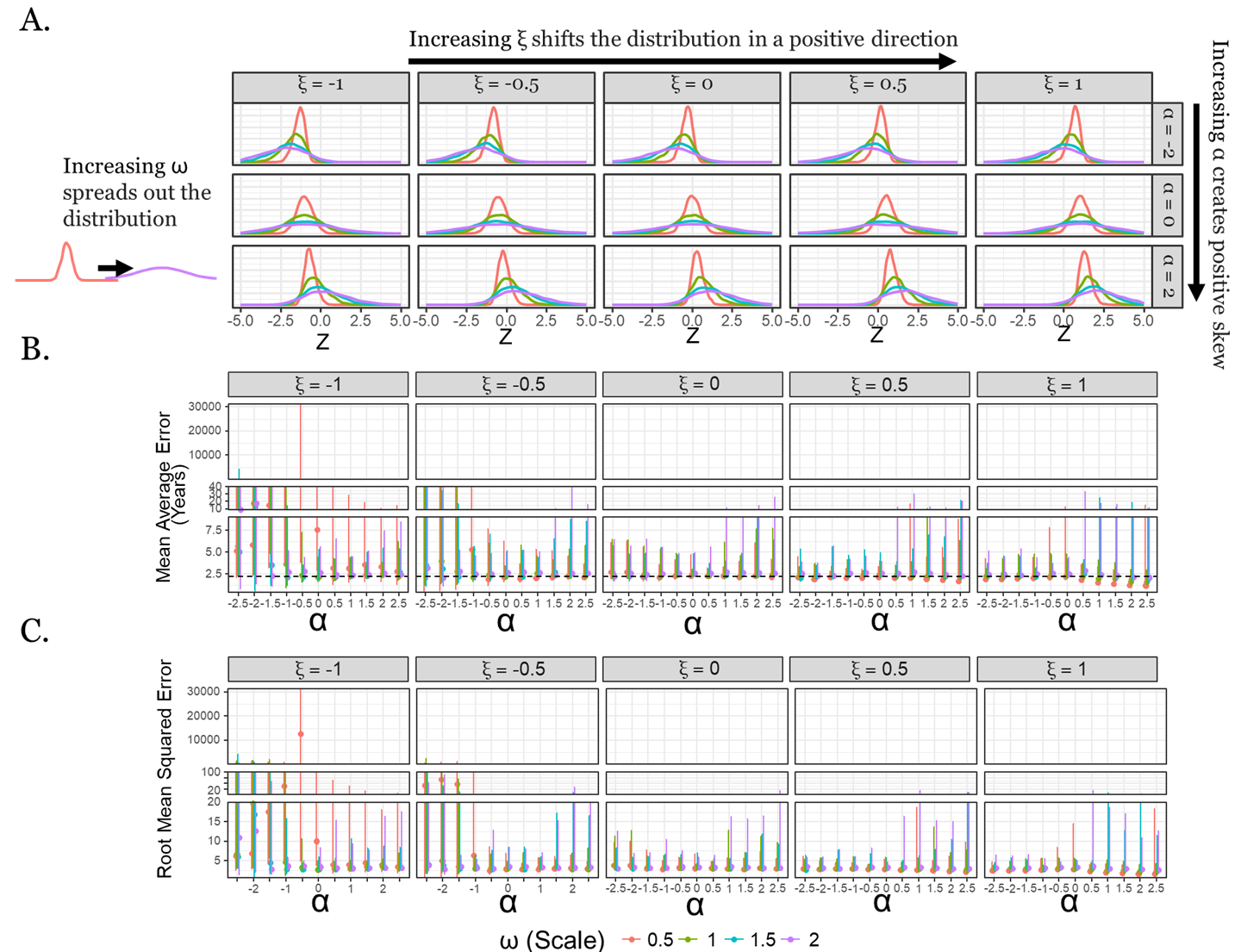

**Supplemental Figure 2.** We generated a range of synthetic datasets characterized by differences in the skew-normal distribution of annualized rates of change values (A). DPM performance across varied skew-normal parameters describing between-individual variability. Lower Mean Average Error (MAE) (B) and Root Mean Squared Error (RMSE) (C) are observed with higher ξ (location), lower ω (scale), and higher α (shape) parameters. The dashed horizontal line represents the mean sampling frequency in the MAE plot (B). Parameter sets characterized by low ξ, low ω, and low α yield errors that exceed the expected sampling variance.

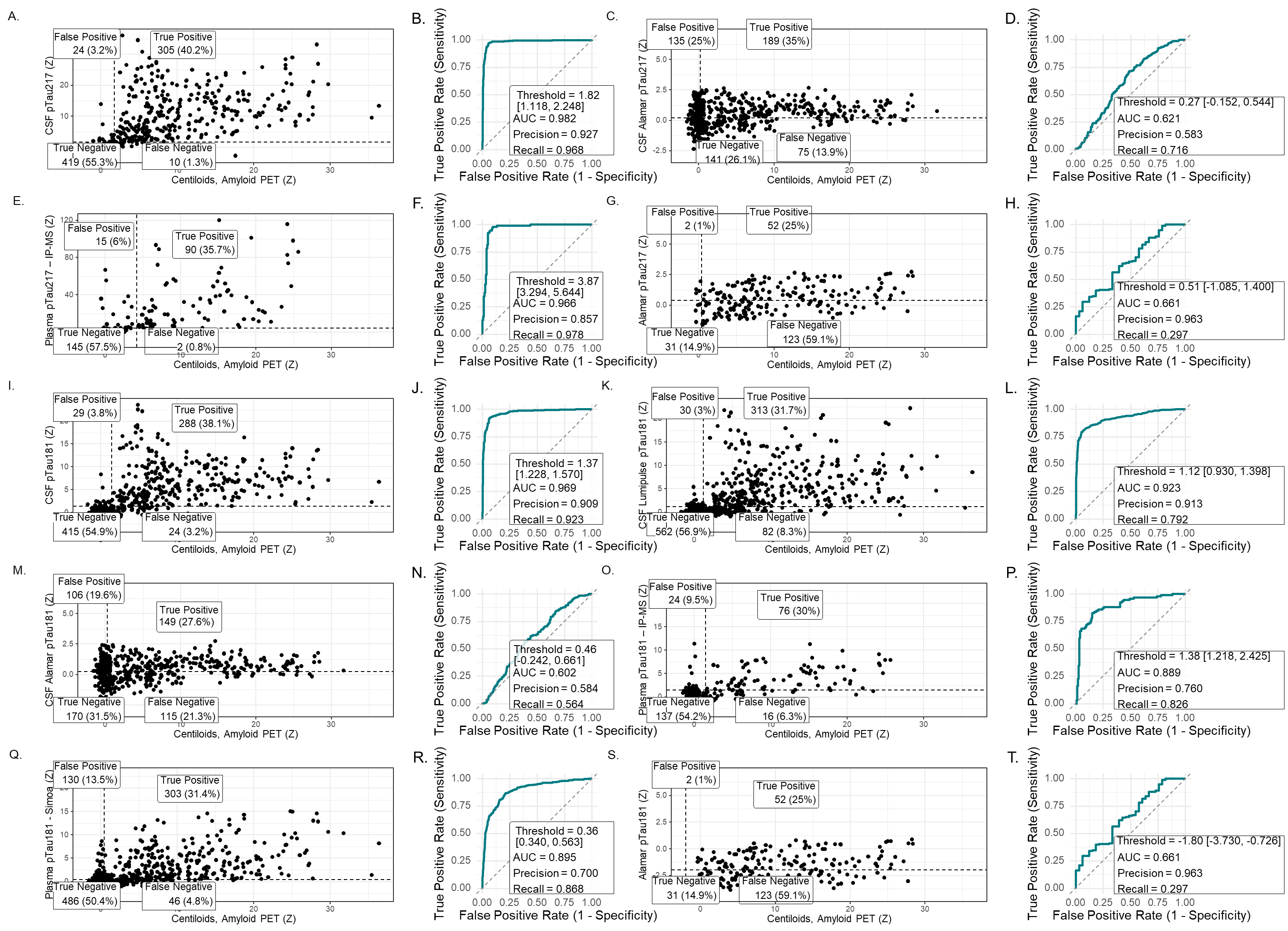

**Supplemental Figure 3.** All fluid biomarkers considered have relatively high correlations with amyloid PET (A, C, E, G, I, K, M, O, Q, S). Dashed lines indicate thresholds for amyloid positivity. The threshold for amyloid PET positivity is literature driven (18 CL, 2.60 Z), while the threshold for biomarker positivity is derived from a cutpoint analysis and varies by biomarker. Corresponding ROC curves (B, D, F, H, J, L, N, P, R, T) summarize classification performance.

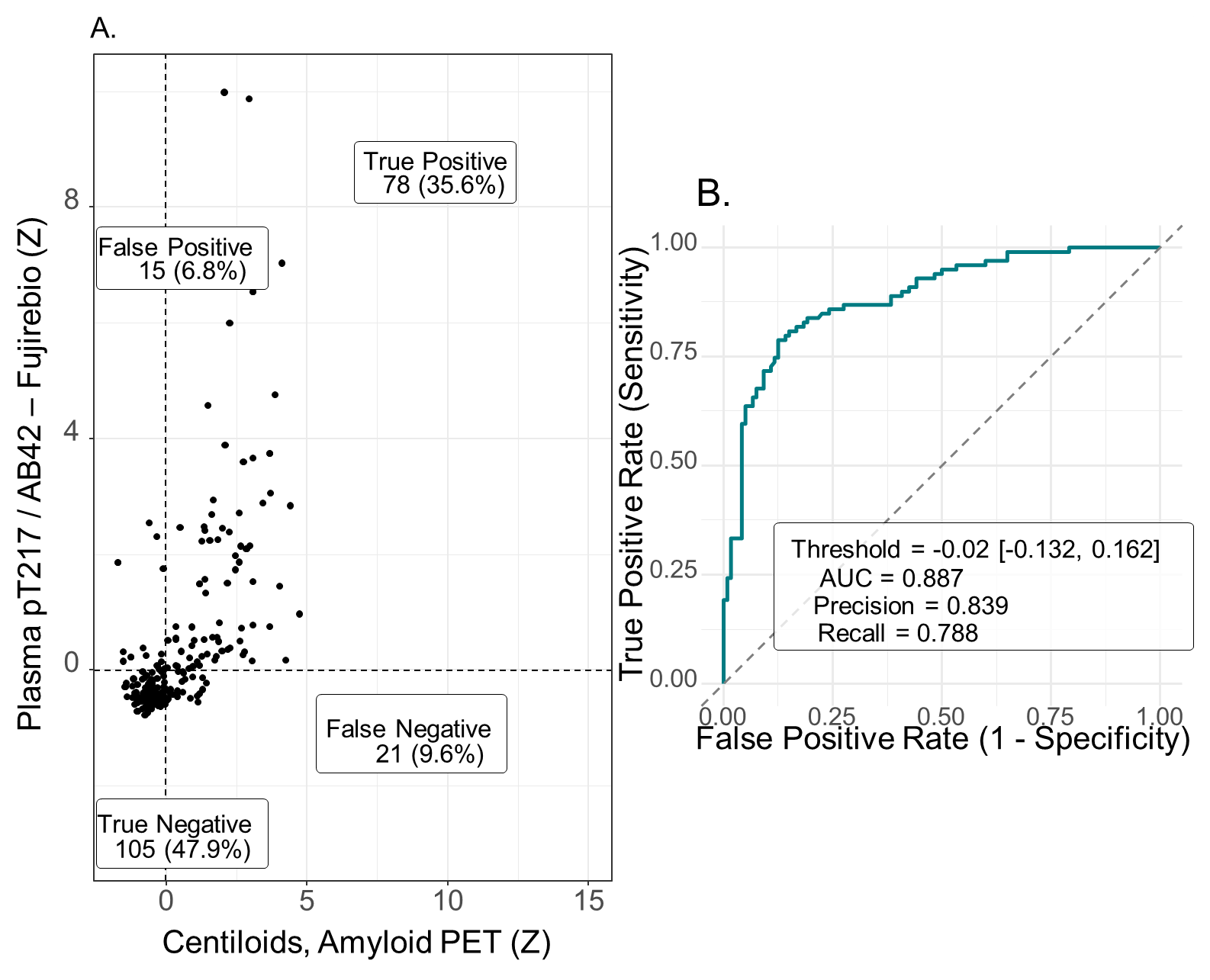

**Supplemental Figure 4.** In ADNI, plasma pTau217/AB42 has good concordance with amyloid PET (A). Dashed lines indicate thresholds for amyloid positivity. The threshold for amyloid PET positivity is literature driven (18 CL, -0.151 Z), while the threshold for biomarker positivity is derived from a cutpoint analysis with the result of 0.00648 pg/mL, 0.0258 Z. The corresponding ROC curves summarizes classification performance (B).

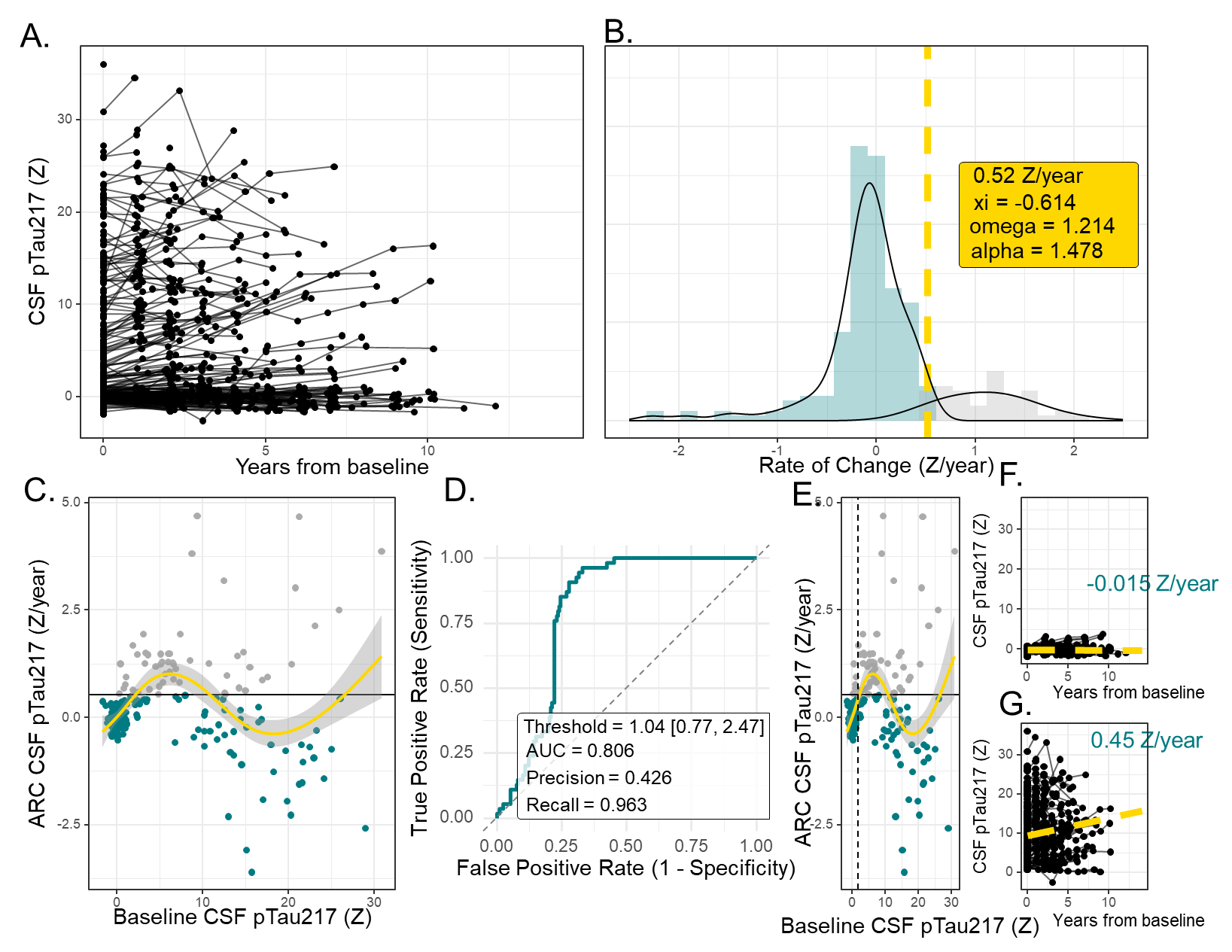

**Supplemental Figure 5. Characterization and Data Selection Methods associated with CSF pTau217 [IP-MS].**

Longitudinal CSF pTau217 (Z-scores) for all participants, plotted relative to years from baseline (A). Distribution of annualized rates of change with fitted skew-normal and Gaussian-mixture components used to identify above-normal accumulation (B). Relationship between annualized rate of change and baseline biomarker level, highlighting individuals exceeding the rate threshold (C). Cutpoint analysis identifying the optimal baseline value for predicting above-normal accumulation (D). Individuals meeting the new criteria for reliable accumulation are shown to the right of the dashed line (E). Individual trajectories from individuals below (F) and above (G) the baseline threshold illustrate unstable versus reliable accumulation patterns. Only participants with baseline biomarker level above the threshold for reliable accumulation were included in downstream disease-progression modeling.

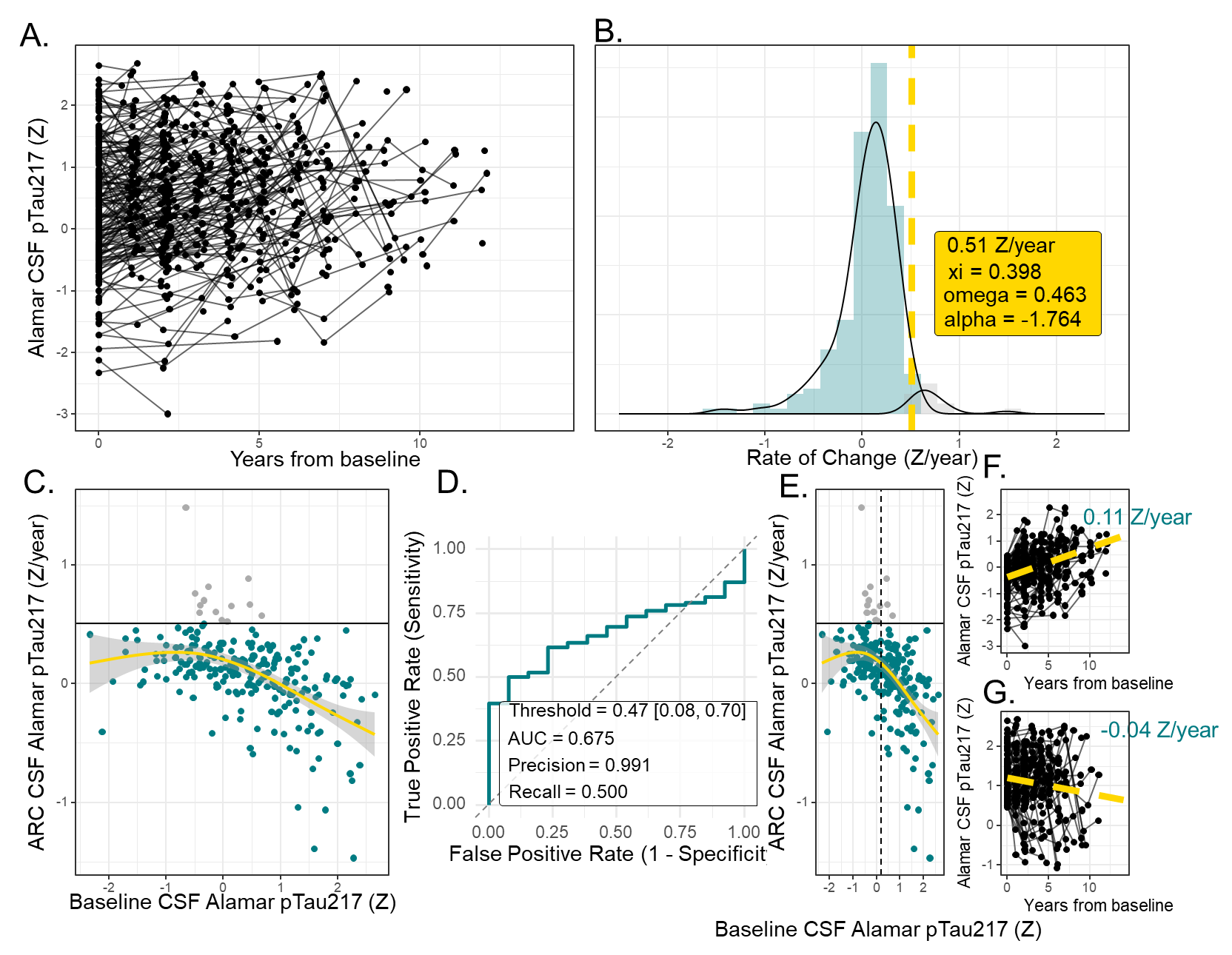

**Supplemental Figure 6. Characterization and Data Selection Methods associated with CSF pTau217 [Alamar NULISA].**

Longitudinal CSF pTau217 (Z-scores) for all participants, plotted relative to years from baseline (A). Distribution of annualized rates of change with fitted skew-normal and Gaussian-mixture components used to identify above-normal accumulation (B). Relationship between annualized rate of change and baseline biomarker level, highlighting individuals exceeding the rate threshold (C). Cutpoint analysis identifying the optimal baseline value for predicting above-normal accumulation (D). Individuals meeting the new criteria for reliable accumulation are shown to the right of the dashed line (E). Individual trajectories from individuals below (F) and above (G) the baseline threshold illustrate unstable versus reliable accumulation patterns. Only participants with baseline biomarker level above the threshold for reliable accumulation were included in downstream disease-progression modeling.

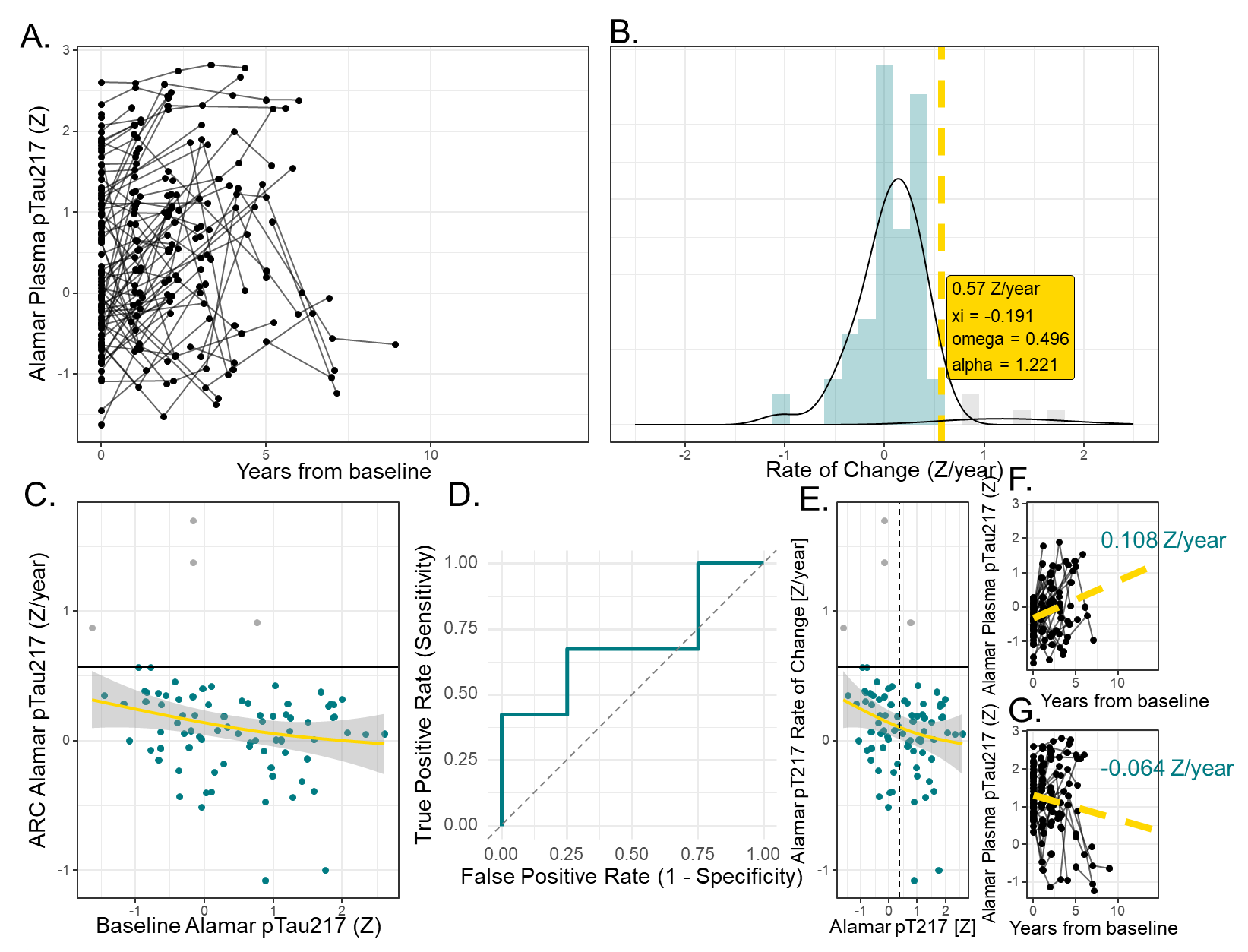

**Supplemental Figure 7. Characterization and Data Selection Methods associated with Plasma pTau217 [Alamar NULISA].**

Longitudinal Plasma pTau217 (Z-scores) for all participants, plotted relative to years from baseline (A). Distribution of annualized rates of change with fitted skew-normal and Gaussian-mixture components used to identify above-normal accumulation (B). Relationship between annualized rate of change and baseline biomarker level, highlighting individuals exceeding the rate threshold (C). Cutpoint analysis identifying the optimal baseline value for predicting above-normal accumulation (D). Individuals meeting the new criteria for reliable accumulation are shown to the right of the dashed line (E). Individual trajectories from individuals below (F) and above (G) the baseline threshold illustrate unstable versus reliable accumulation patterns. Only participants with baseline biomarker level above the threshold for reliable accumulation were included in downstream disease-progression modeling.

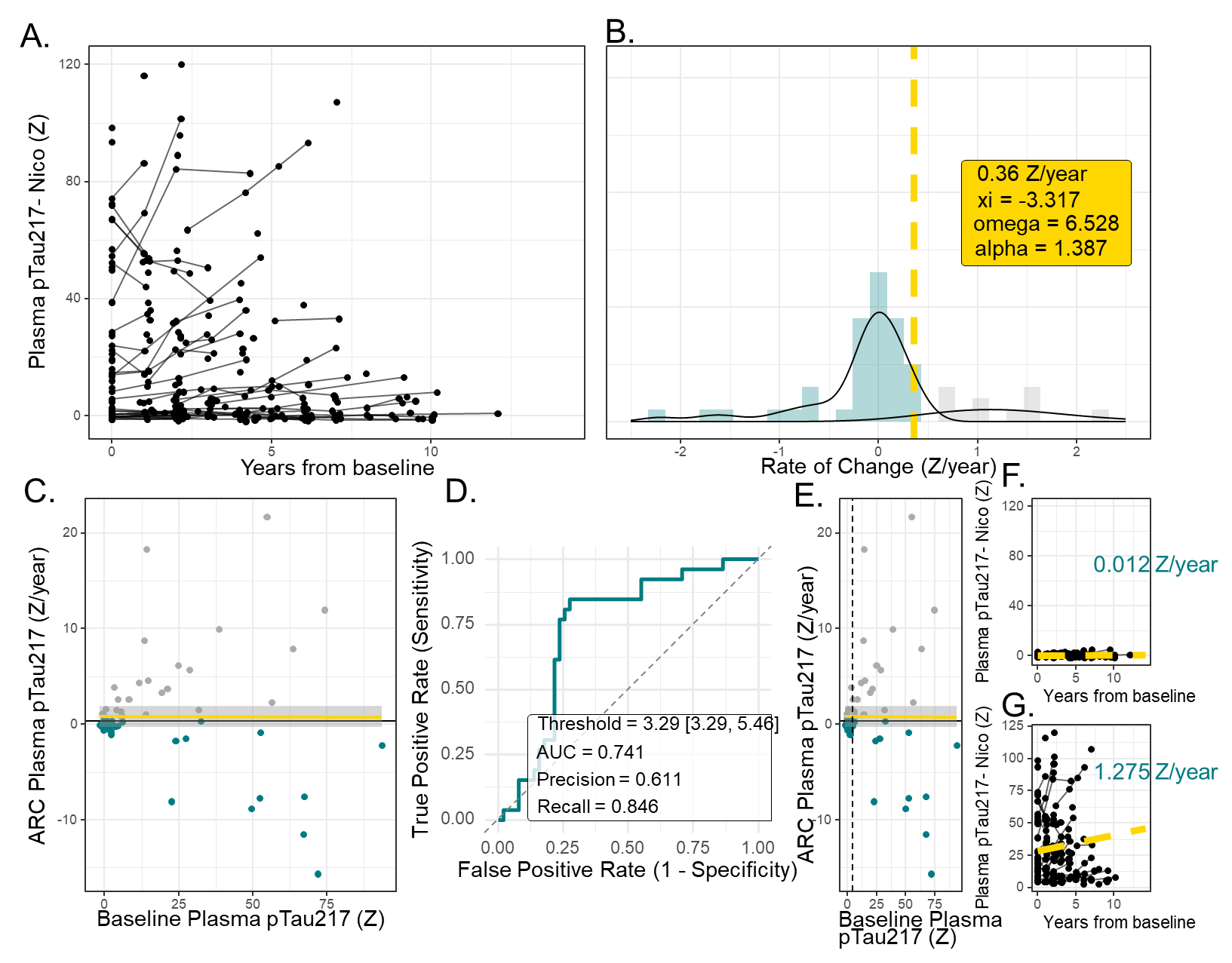
 **Supplemental Figure 8. Characterization and Data Selection Methods associated with Plasma pTau217 [IP-MS].**

Longitudinal Plasma pTau217 (Z-scores) for all participants, plotted relative to years from baseline (A). Distribution of annualized rates of change with fitted skew-normal and Gaussian-mixture components used to identify above-normal accumulation (B). Relationship between annualized rate of change and baseline biomarker level, highlighting individuals exceeding the rate threshold (C). Cutpoint analysis identifying the optimal baseline value for predicting above-normal accumulation (D). Individuals meeting the new criteria for reliable accumulation are shown to the right of the dashed line (E). Individual trajectories from individuals below (F) and above (G) the baseline threshold illustrate unstable versus reliable accumulation patterns. Only participants with baseline biomarker level above the threshold for reliable accumulation were included in downstream disease-progression modeling.

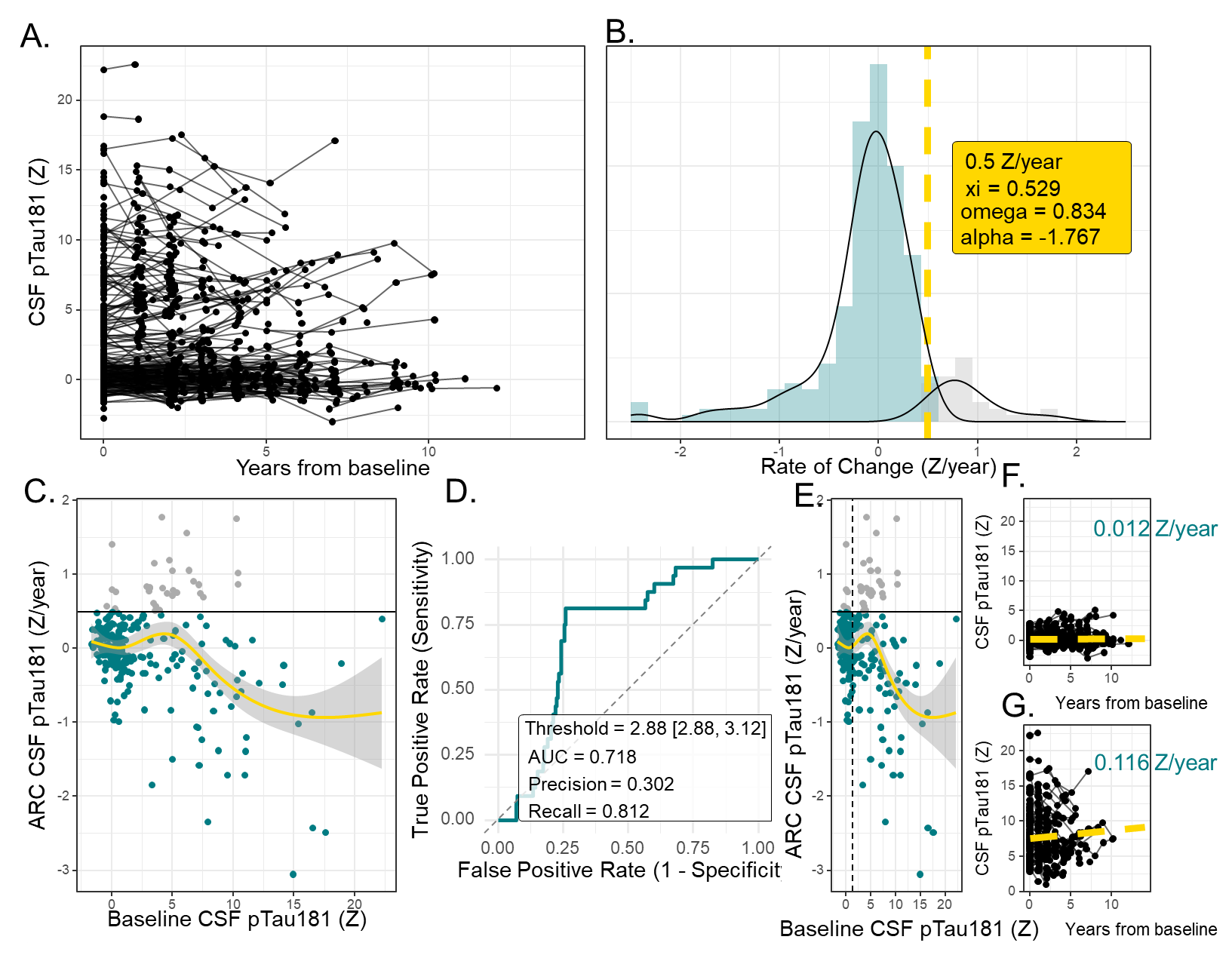

**Supplemental Figure 9. Characterization and Data Selection Methods associated with CSF pTau181 [IP-MS].**

Longitudinal CSF pTau181 (Z-scores) for all participants, plotted relative to years from baseline (A). Distribution of annualized rates of change with fitted skew-normal and Gaussian-mixture components used to identify above-normal accumulation (B). Relationship between annualized rate of change and baseline biomarker level, highlighting individuals exceeding the rate threshold (C). Cutpoint analysis identifying the optimal baseline value for predicting above-normal accumulation (D). Individuals meeting the new criteria for reliable accumulation are shown to the right of the dashed line (E). Individual trajectories from individuals below (F) and above (G) the baseline threshold illustrate unstable versus reliable accumulation patterns. Only participants with baseline biomarker level above the threshold for reliable accumulation were included in downstream disease-progression modeling.

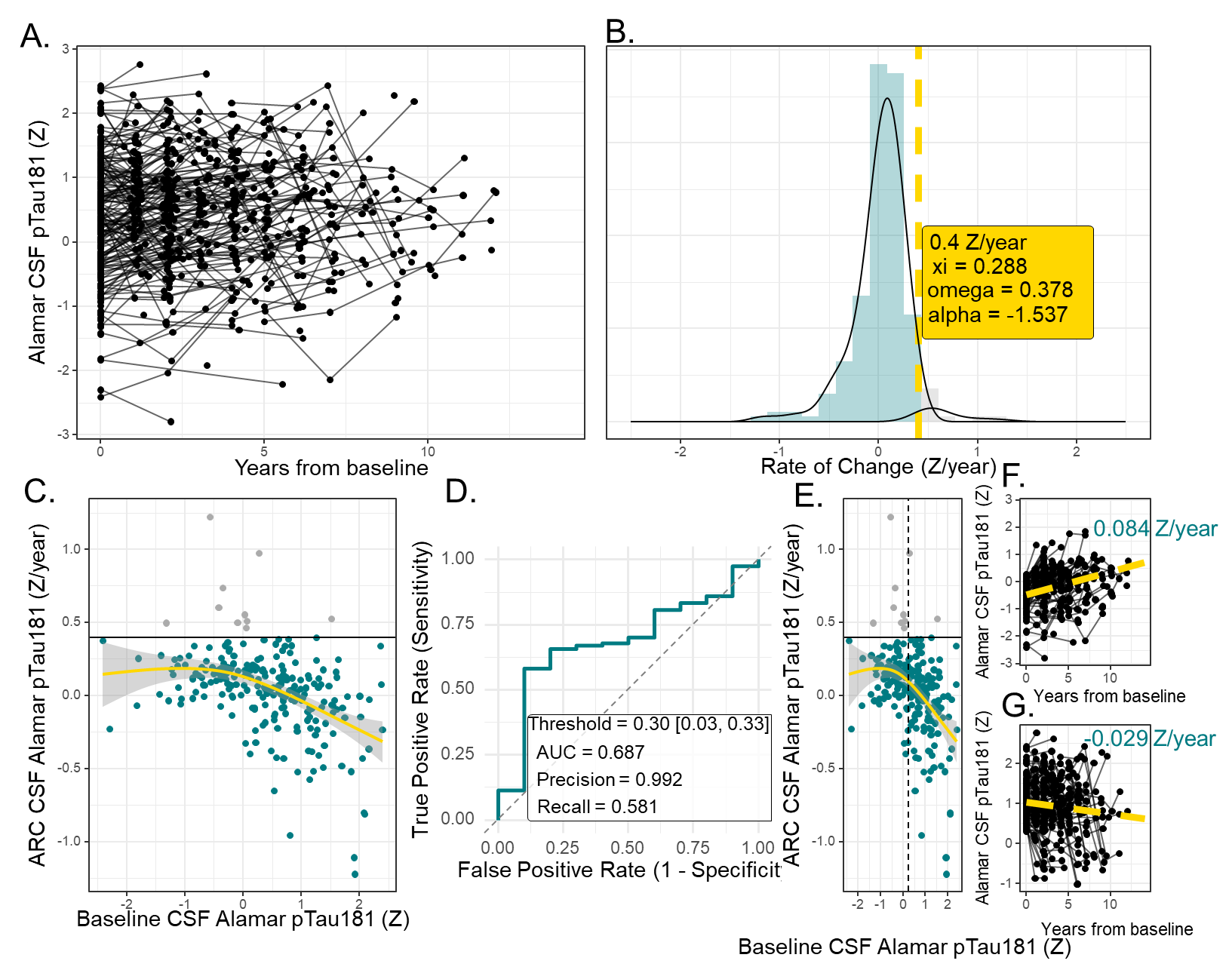

**Supplemental Figure 10. Characterization and Data Selection Methods associated with CSF pTau181 [Alamar NULISA].**

Longitudinal CSF pTau181 (Z-scores) for all participants, plotted relative to years from baseline (A). Distribution of annualized rates of change with fitted skew-normal and Gaussian-mixture components used to identify above-normal accumulation (B). Relationship between annualized rate of change and baseline biomarker level, highlighting individuals exceeding the rate threshold (C). Cutpoint analysis identifying the optimal baseline value for predicting above-normal accumulation (D). Individuals meeting the new criteria for reliable accumulation are shown to the right of the dashed line (E). Individual trajectories from individuals below (F) and above (G) the baseline threshold illustrate unstable versus reliable accumulation patterns. Only participants with baseline biomarker level above the threshold for reliable accumulation were included in downstream disease-progression modeling.

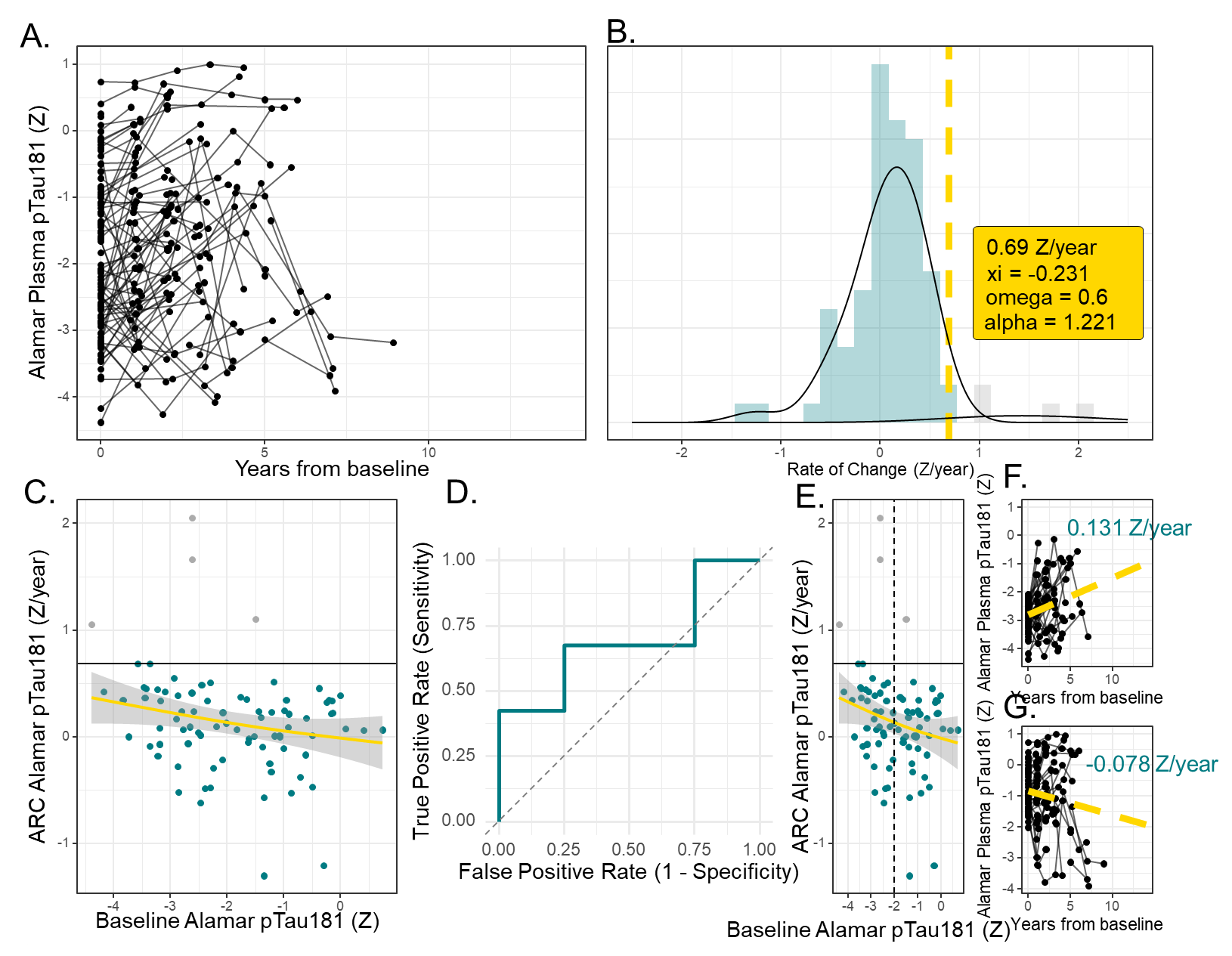

**Supplemental Figure 11. Characterization and Data Selection Methods associated with Plasma pTau181 [Alamar NULISA].**

Longitudinal Plasma pTau181 (Z-scores) for all participants, plotted relative to years from baseline (A). Distribution of annualized rates of change with fitted skew-normal and Gaussian-mixture components used to identify above-normal accumulation (B). Relationship between annualized rate of change and baseline biomarker level, highlighting individuals exceeding the rate threshold (C). Cutpoint analysis identifying the optimal baseline value for predicting above-normal accumulation (D). Individuals meeting the new criteria for reliable accumulation are shown to the right of the dashed line (E). Individual trajectories from individuals below (F) and above (G) the baseline threshold illustrate unstable versus reliable accumulation patterns. Only participants with baseline biomarker level above the threshold for reliable accumulation were included in downstream disease-progression modeling.

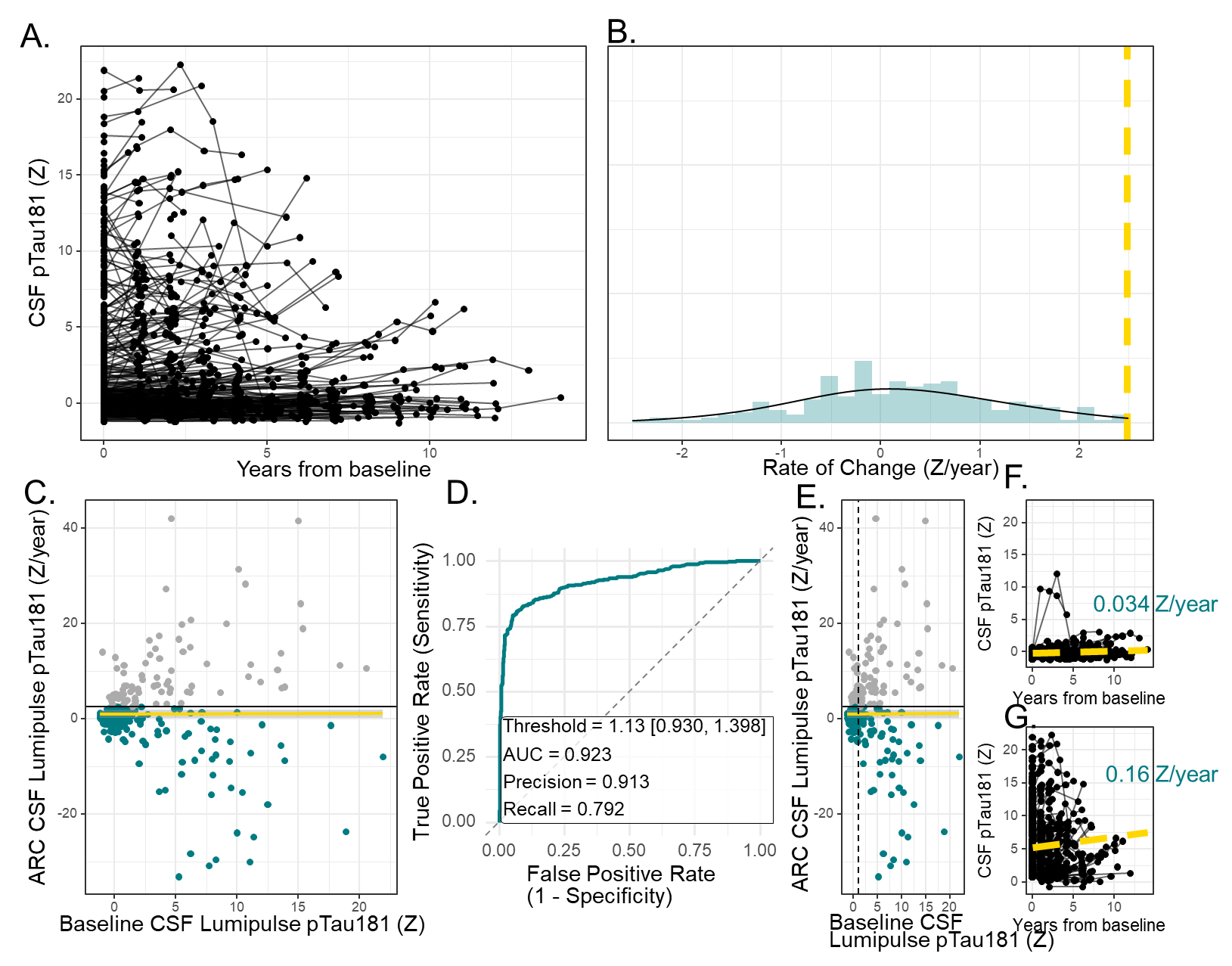

**Supplemental Figure 12. Characterization and Data Selection Methods associated with CSF pTau181 [Lumipulse].**

Longitudinal CSF pTau181 (Z-scores) for all participants, plotted relative to years from baseline (A). Distribution of annualized rates of change with fitted skew-normal and Gaussian-mixture components used to identify above-normal accumulation (B). Relationship between annualized rate of change and baseline biomarker level, highlighting individuals exceeding the rate threshold (C). Cutpoint analysis identifying the optimal baseline value for predicting above-normal accumulation (D). Individuals meeting the new criteria for reliable accumulation are shown to the right of the dashed line (E). Individual trajectories from individuals below (F) and above (G) the baseline threshold illustrate unstable versus reliable accumulation patterns. Only participants with baseline biomarker level above the threshold for reliable accumulation were included in downstream disease-progression modeling.

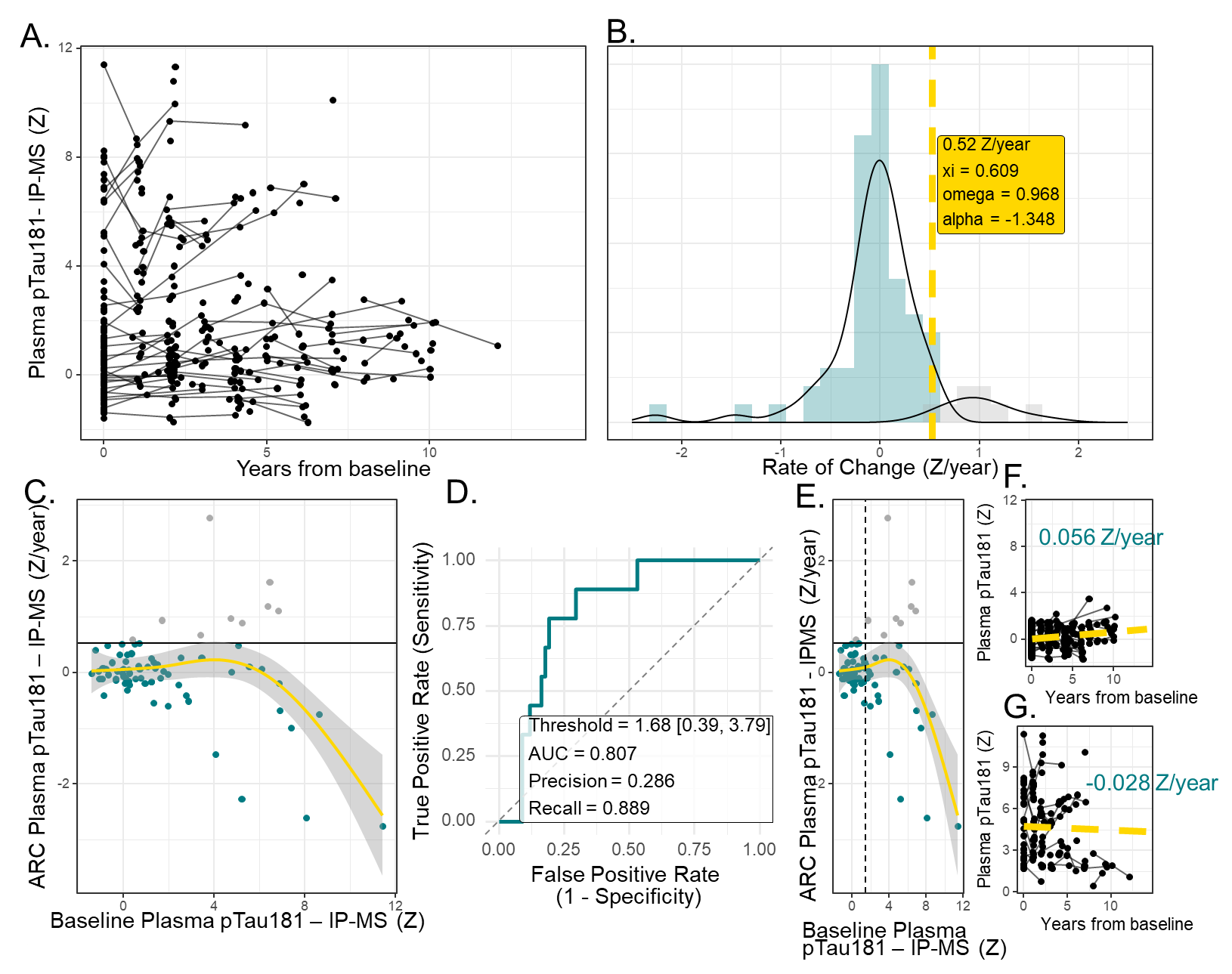

**Supplemental Figure 13. Characterization and Data Selection Methods associated with Plasma pTau181 [IP-MS].**

Longitudinal plasma pTau181 (Z-scores) for all participants, plotted relative to years from baseline (A). Distribution of annualized rates of change with fitted skew-normal and Gaussian-mixture components used to identify above-normal accumulation (B). Relationship between annualized rate of change and baseline biomarker level, highlighting individuals exceeding the rate threshold (C). Cutpoint analysis identifying the optimal baseline value for predicting above-normal accumulation (D). Individuals meeting the new criteria for reliable accumulation are shown to the right of the dashed line (E). Individual trajectories from individuals below (F) and above (G) the baseline threshold illustrate unstable versus reliable accumulation patterns. Only participants with baseline biomarker level above the threshold for reliable accumulation were included in downstream disease-progression modeling.

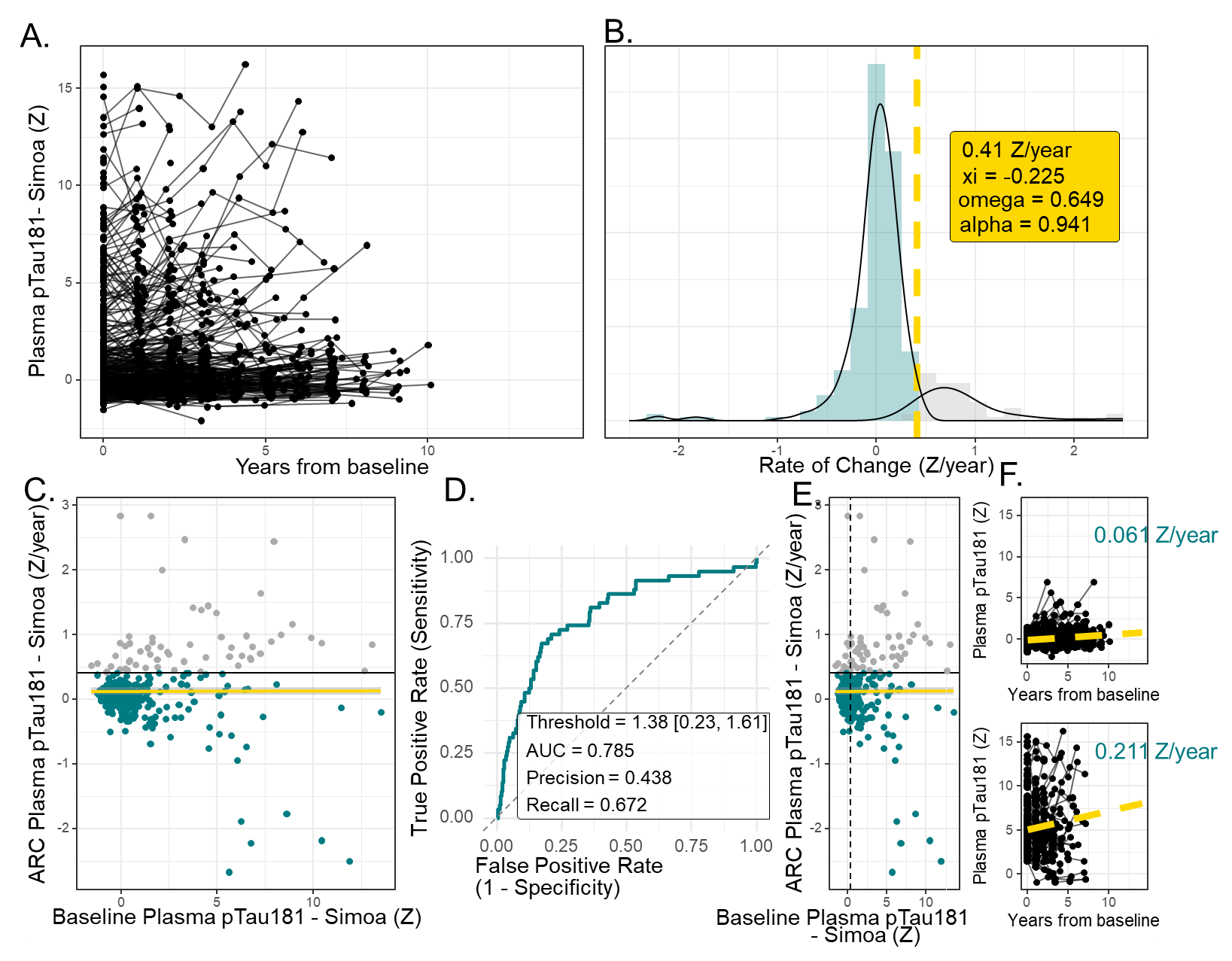

**Supplemental Figure 14. Characterization and Data Selection Methods associated with Plasma pTau181 [Simoa].**

Longitudinal plasma pTau181 (Z-scores) for all participants, plotted relative to years from baseline (A). Distribution of annualized rates of change with fitted skew-normal and Gaussian-mixture components used to identify above-normal accumulation (B). Relationship between annualized rate of change and baseline biomarker level, highlighting individuals exceeding the rate threshold (C). Cutpoint analysis identifying the optimal baseline value for predicting above-normal accumulation (D). Individuals meeting the new criteria for reliable accumulation are shown to the right of the dashed line (E). Individual trajectories from individuals below (F) and above (G) the baseline threshold illustrate unstable versus reliable accumulation patterns. Only participants with baseline biomarker level above the threshold for reliable accumulation were included in downstream disease-progression modeling.

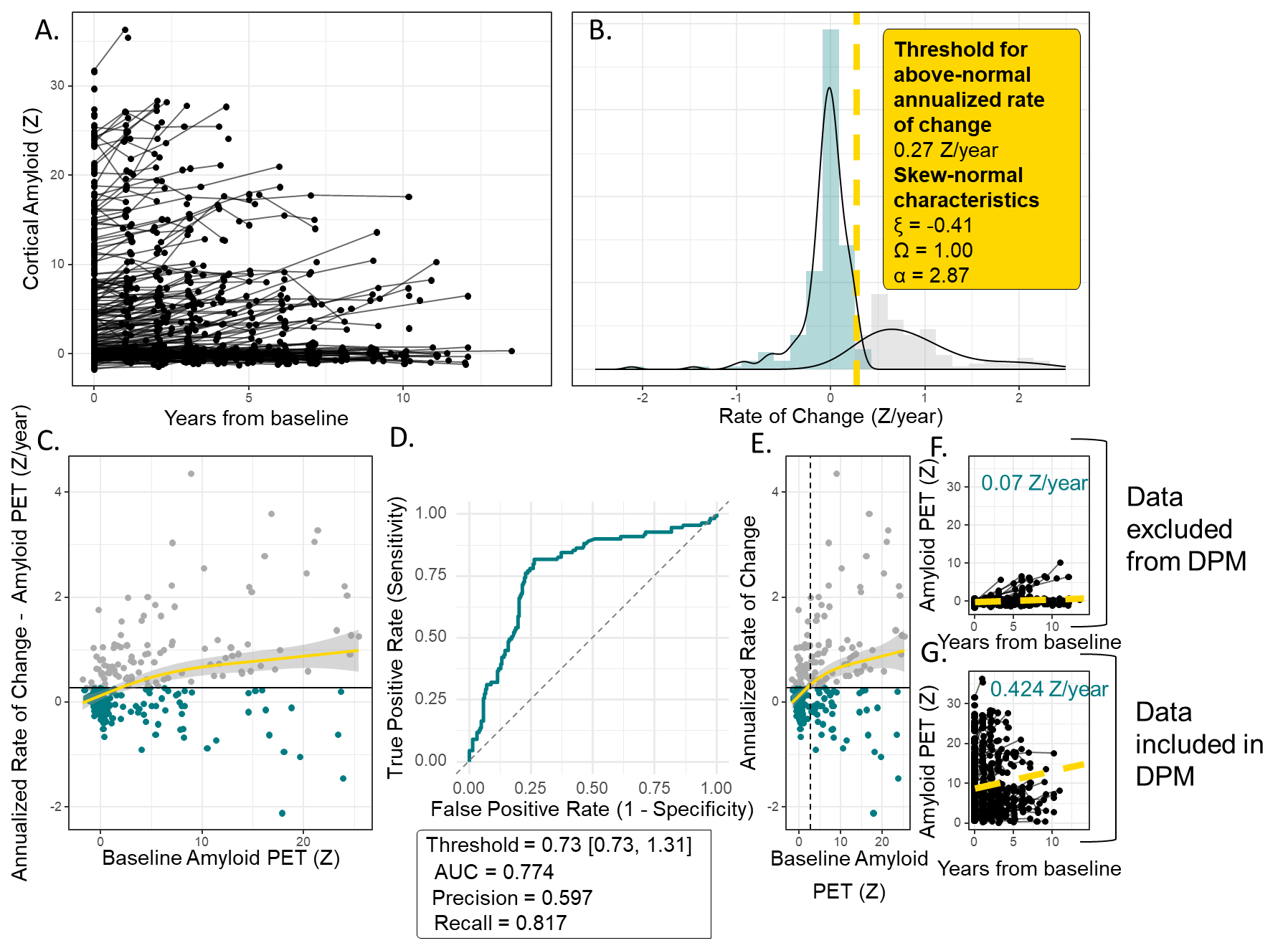

**Supplemental Figure 15. Characterization and Data Selection Methods associated with Amyloid PET, DIAN.**

Longitudinal cortical amyloid PET trajectories (Z-scores) for all participants, plotted relative to years from baseline (A). Distribution of annualized rates of change with fitted skew-normal and Gaussian-mixture components used to identify above-normal accumulation (B). Relationship between annualized rate of change and baseline amyloid PET, highlighting individuals exceeding the rate threshold (C). Cutpoint analysis identifying the optimal baseline value for predicting above-normal accumulation (D). Individuals meeting the new criteria for reliable accumulation are shown to the right of the dashed line (E). Individual trajectories from individuals below (F) and above (G) the baseline threshold illustrate unstable versus reliable accumulation patterns. Only participants with baseline cortical amyloid PET ≥ 0.73 Z were included in downstream disease-progression modeling.

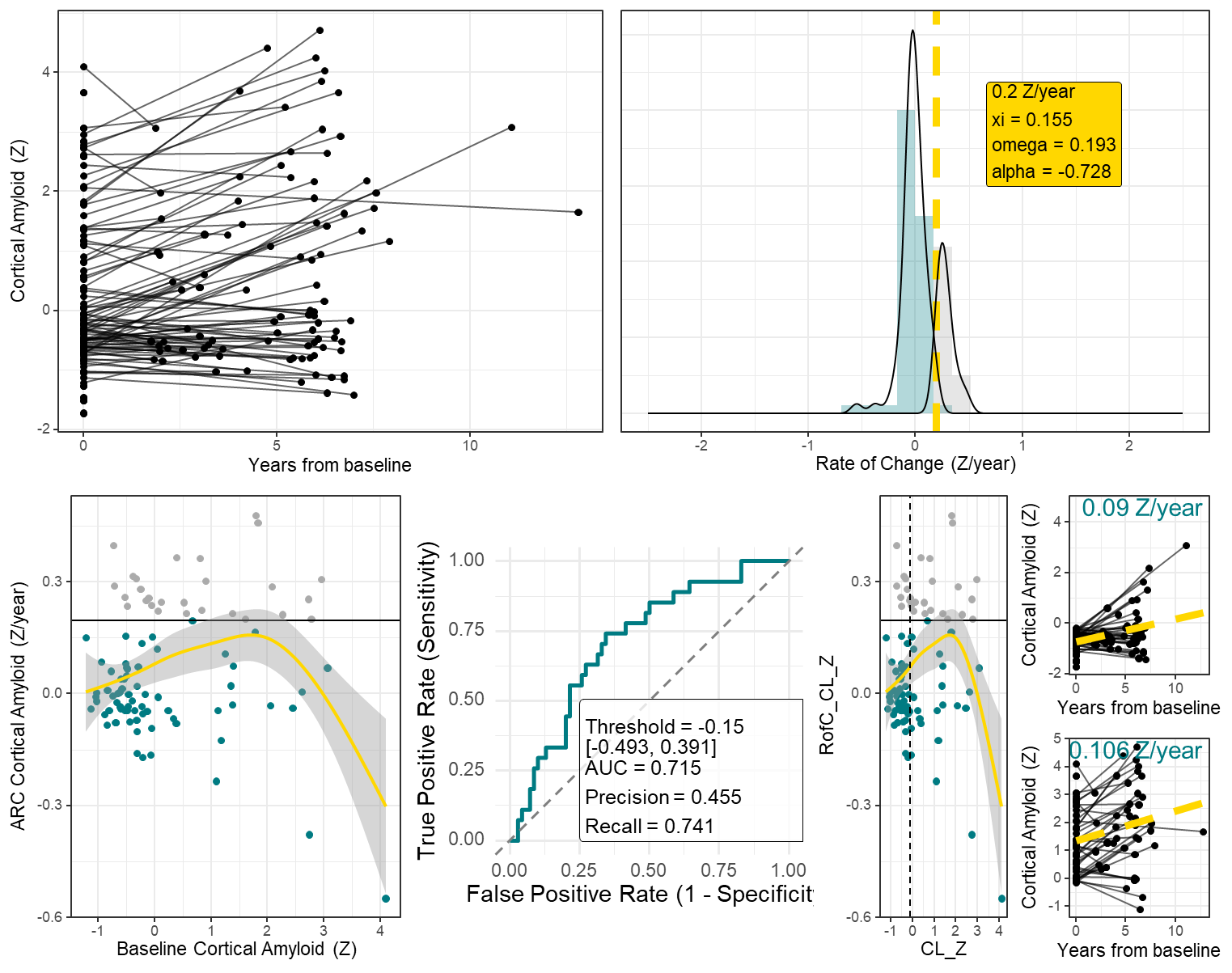

**Supplemental Figure 16. Characterization and Data Selection Methods associated with Amyloid PET, ADNI.**

Longitudinal cortical amyloid PET trajectories (Z-scores) for all participants, plotted relative to years from baseline (A). Distribution of annualized rates of change with fitted skew-normal and Gaussian-mixture components used to identify above-normal accumulation (B). Relationship between annualized rate of change and baseline amyloid PET, highlighting individuals exceeding the rate threshold (C). Cutpoint analysis identifying the optimal baseline value for predicting above-normal accumulation (D). Individuals meeting the new criteria for reliable accumulation are shown to the right of the dashed line (E). Individual trajectories from individuals below (F) and above (G) the baseline threshold illustrate unstable versus reliable accumulation patterns.

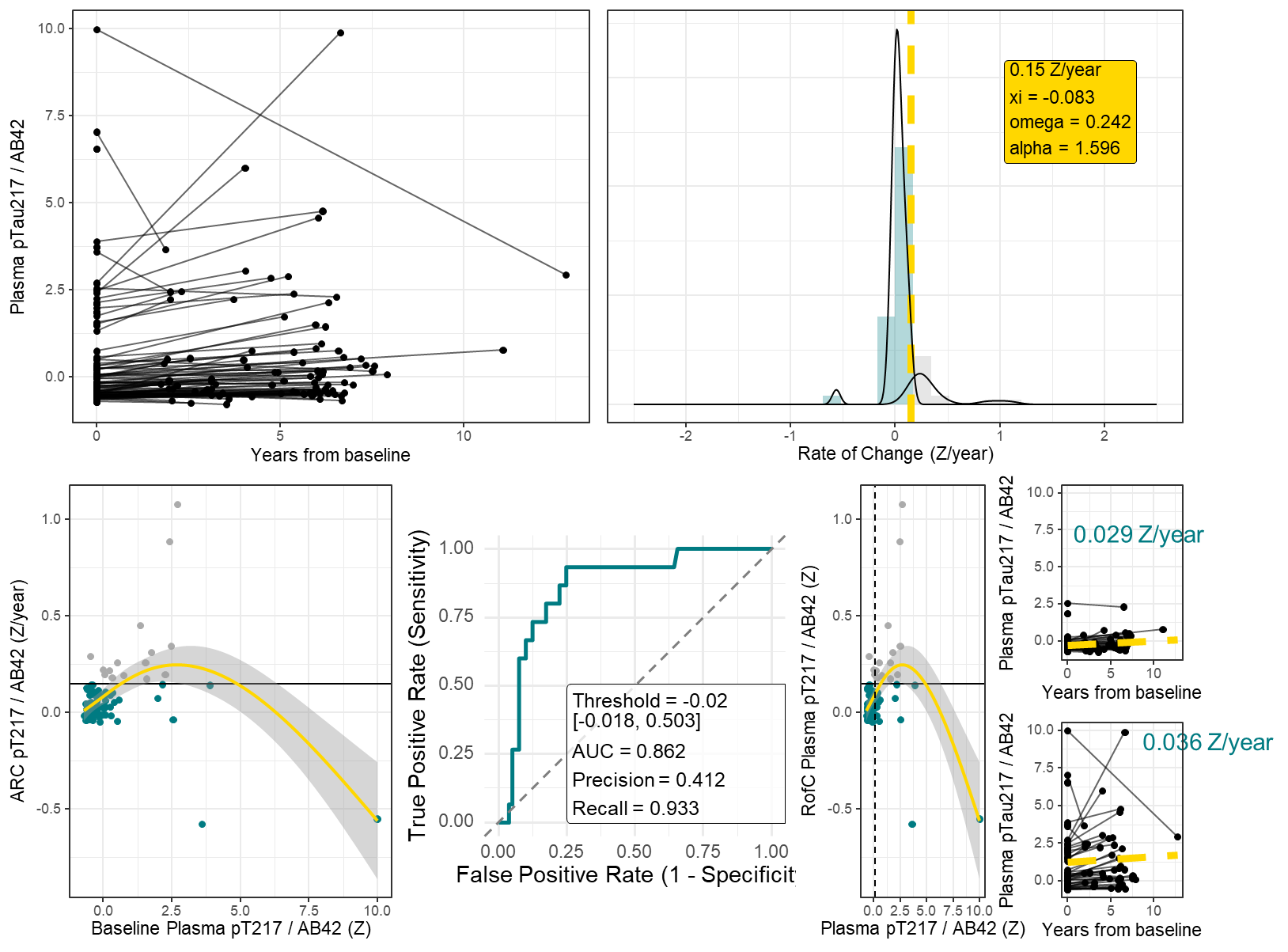

**Supplemental Figure 17. Characterization and Data Selection Methods associated with Plasma pTau217 / AB42 - Fujirebio, ADNI.**

Longitudinal plasma pTau217 / AB42 trajectories (Z-scores) for all participants, plotted relative to years from baseline (A). Distribution of annualized rates of change with fitted skew-normal and Gaussian-mixture components used to identify above-normal accumulation (B). Relationship between annualized rate of change and baseline plasma, highlighting individuals exceeding the rate threshold (C). Cutpoint analysis identifying the optimal baseline value for predicting above-normal accumulation (D). Individuals meeting the new criteria for reliable accumulation are shown to the right of the dashed line (E). Individual trajectories from individuals below (F) and above (G) the baseline threshold illustrate unstable versus reliable accumulation patterns.

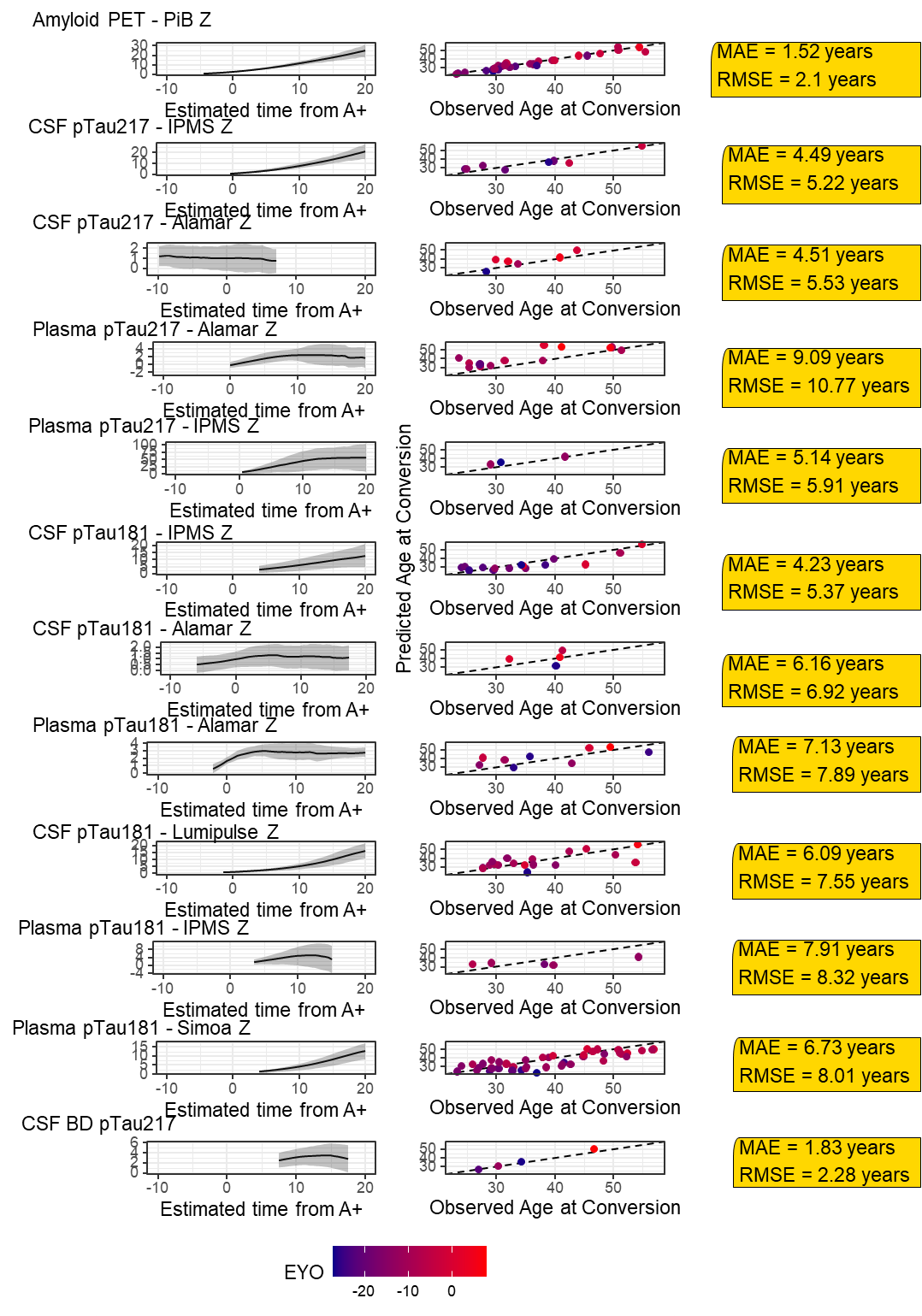

**Supplemental Figure 18. Application of Disease Progression Modeling to DIAN Data.** Model fits for each biomarker are shown (Left Panel). Note that because of minimum thresholds for reliable accumulation, not all biomarkers generate temporal predictions prior to conversion to amyloid positivity (time = 0). These fits were compared to actual observed conversion events from amyloid negative to amyloid positive as were available in the data (Center Panel). We calculated the Mean Average Error (MAE) and Root Mean Squared Error (RMSE) for each evaluated model (Right Panel). When evaluating these parameters, it’s important to keep in mind that some biomarkers like CSF Brain Derived (BD) pT217 and Plasma pT217 – IPMS have very few observed conversion events available for validation.

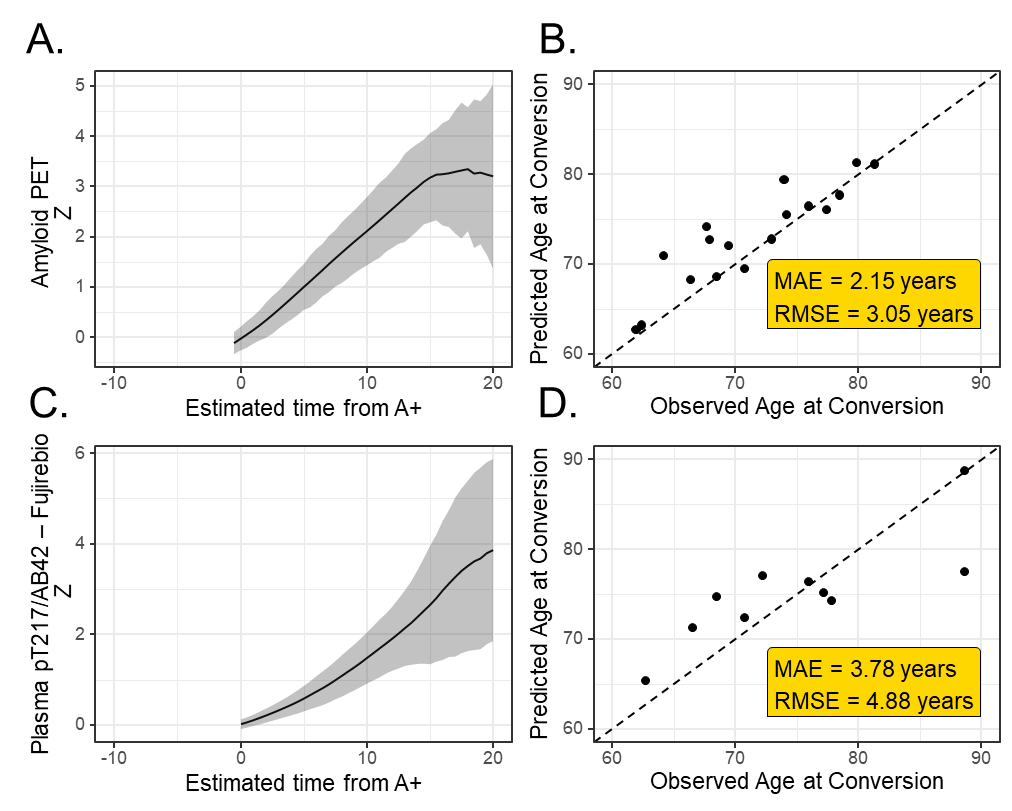

**Supplemental Figure 19. Application of Disease Progression Modeling to ADNI Data.** Model fits for each biomarker are shown (Left Panels). Note that because of minimum thresholds for reliable accumulation, not all biomarkers generate temporal predictions prior to conversion to amyloid positivity (time = 0). These fits were compared to actual observed conversion events from amyloid negative to amyloid positive as were available in the data (Right Panels). We calculated the Mean Average Error (MAE) and Root Mean Squared Error (RMSE) for each evaluated model.

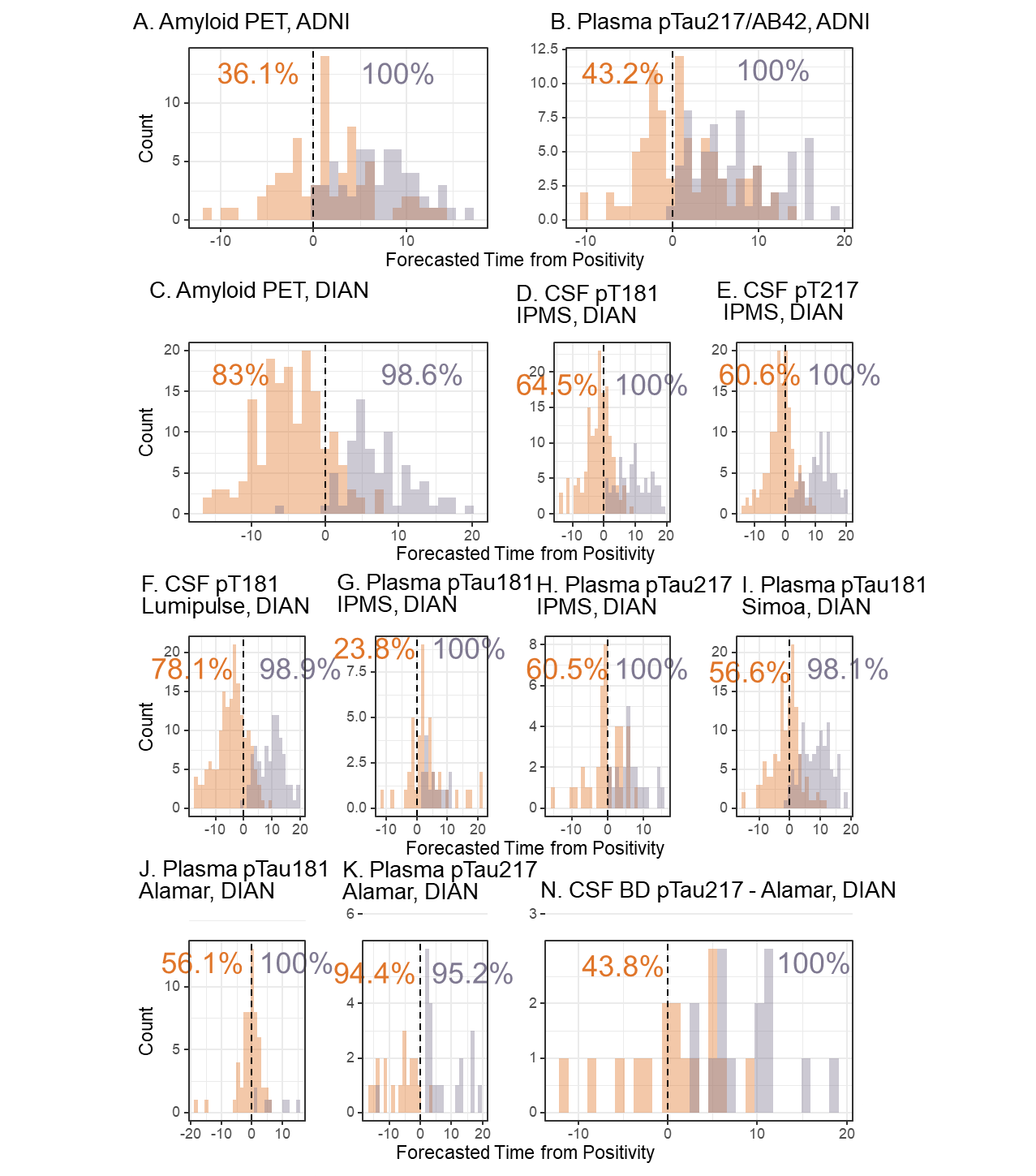

**Supplemental Figure 20.** Because there were a limited number of observed conversion events for each of the biomarkers, we performed an evaluation of the model calibration. For individuals who were amyloid negative throughout enrollment of the study, we estimated their time from positivity based on model fit at their baseline visit. Then we randomly selected one follow up visit for this individual and calculated their estimated time from positivity by summing their original estimated time from positivity and the time elapsed since baseline visit. Theoretically, all of these individuals should have had a negative time estimate, as all of these individuals were, we knew, negative throughout the study. These individuals are shown in orange in each of the above histograms. Model success ranged from as low as 36.1% in ADNI Amyloid PET data, which means that of all of the individuals who were known to be amyloid negative throughout the study, only 36.1% of them were predicted to be negative throughout the study, to values as high as 94.4% in the case of the plasma pTau217 Alamar in DIAN data. Conversely, we identified all individuals who were amyloid positive throughout the study. We estimated their time from positivity based on the model fit for their final visit. Then we randomly selected one prior visit for each individual and calculated their estimated time from positivity by subtracting the time that had elapsed during the study from the original estimated time from positivity. All of these individuals should have had a positive time estimate, as all of these individuals were known to be positive throughout the study and shown in purple on the histograms. Model success ranged from 95.2% to 100%. This calibration analysis suggests that the model performs much better in individuals who are biomarker positive than biomarker negative. This makes sense because individuals who are biomarker negative are much more likely to be below the signal floor, even after the application of the “reliable accumulator” rule, leading to greater stochasticity in longitudinal predictions. *Note that non-brain derived CSF – Alamar data in DIAN is not presented because all individuals exhibited such signal fluctuation as to have both positive and negative visits.*

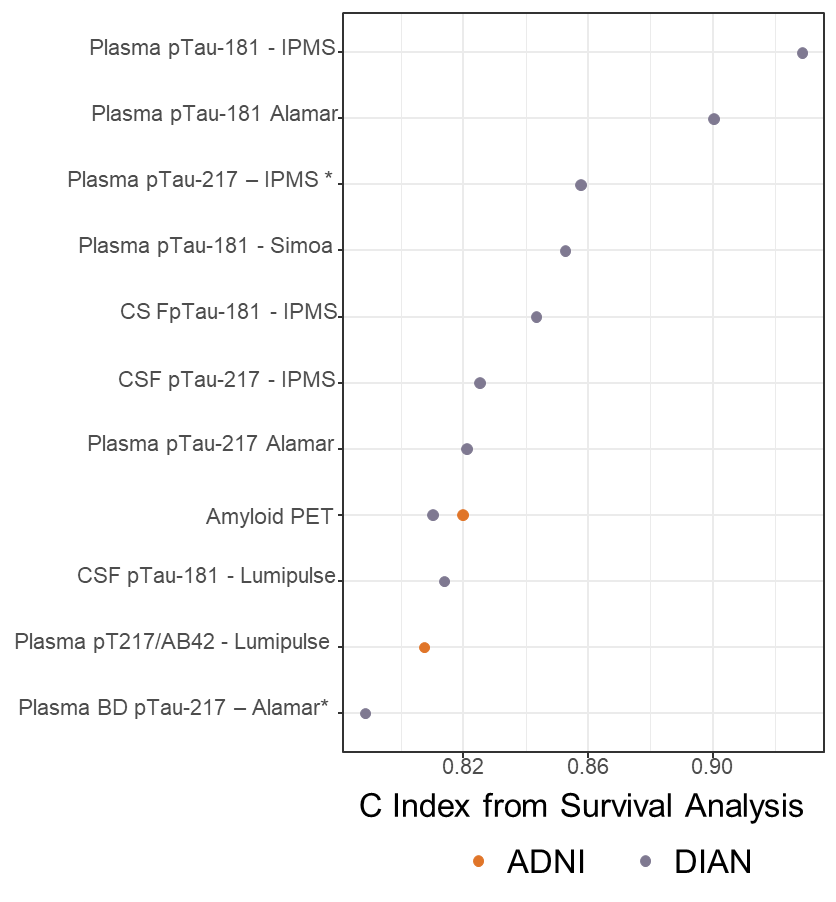

**Supplemental Figure 21.** We performed a survival analysis with subsequent C index calculation to evaluate how well the amyloid time model preserved relative disease ordering. In all cases, the model achieved good performance, with all C indices over 0.80. This suggests that individuals who convert at younger ages are, regardless of biomarker, largely identified as relatively earlier converters by the amyloid time modeling technique. **Biomarkers where 5 or fewer conversion events were observed are indicated with an asterisk. Interpretation of results should be done in a cautious manner.*
