## Supplementary figures and images for "Biomarker Variability Limits Individualized Amyloid Time Estimation in Alzheimer Disease"

### Supp Video - 3% Within Individ Var

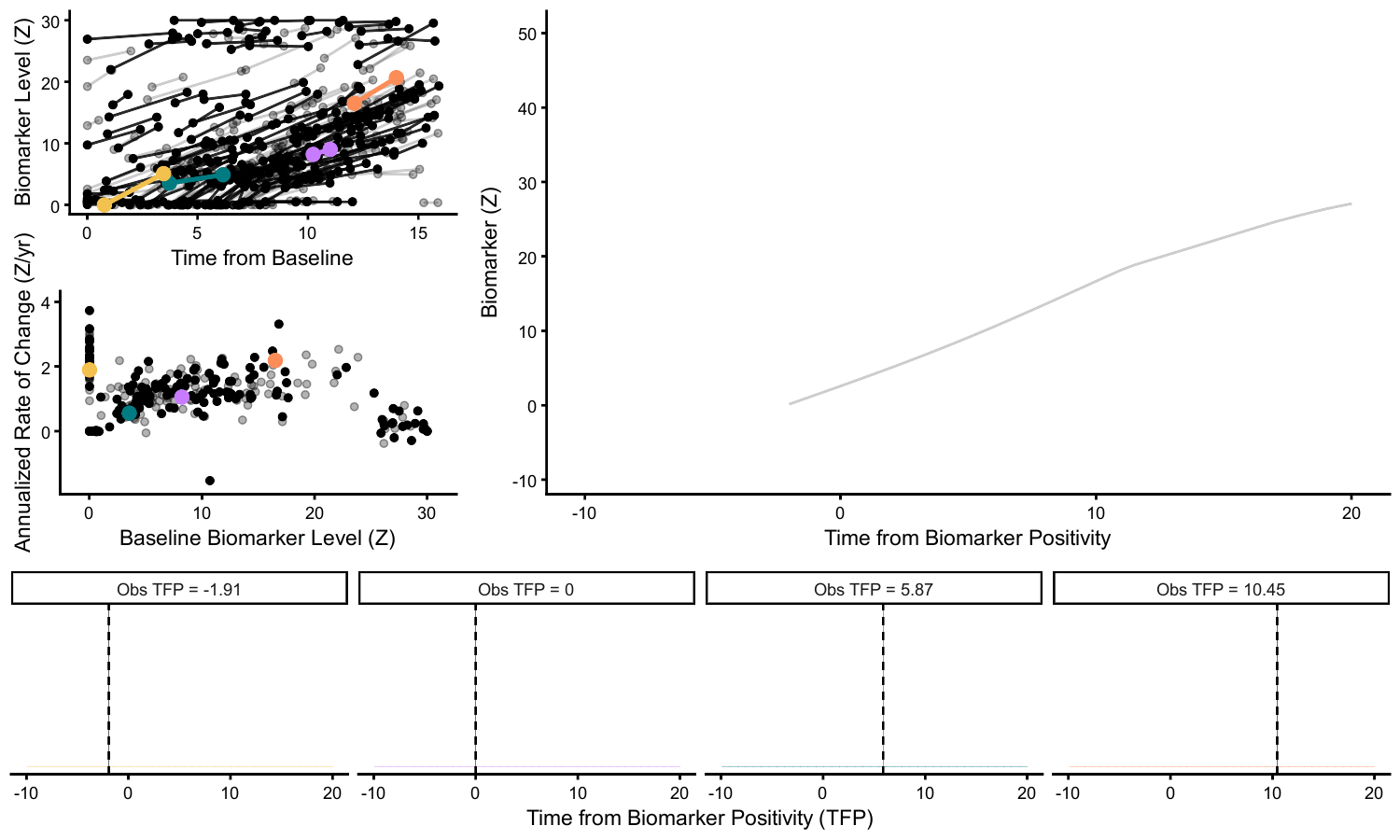

### Supp Video - 20% Within Individ Var

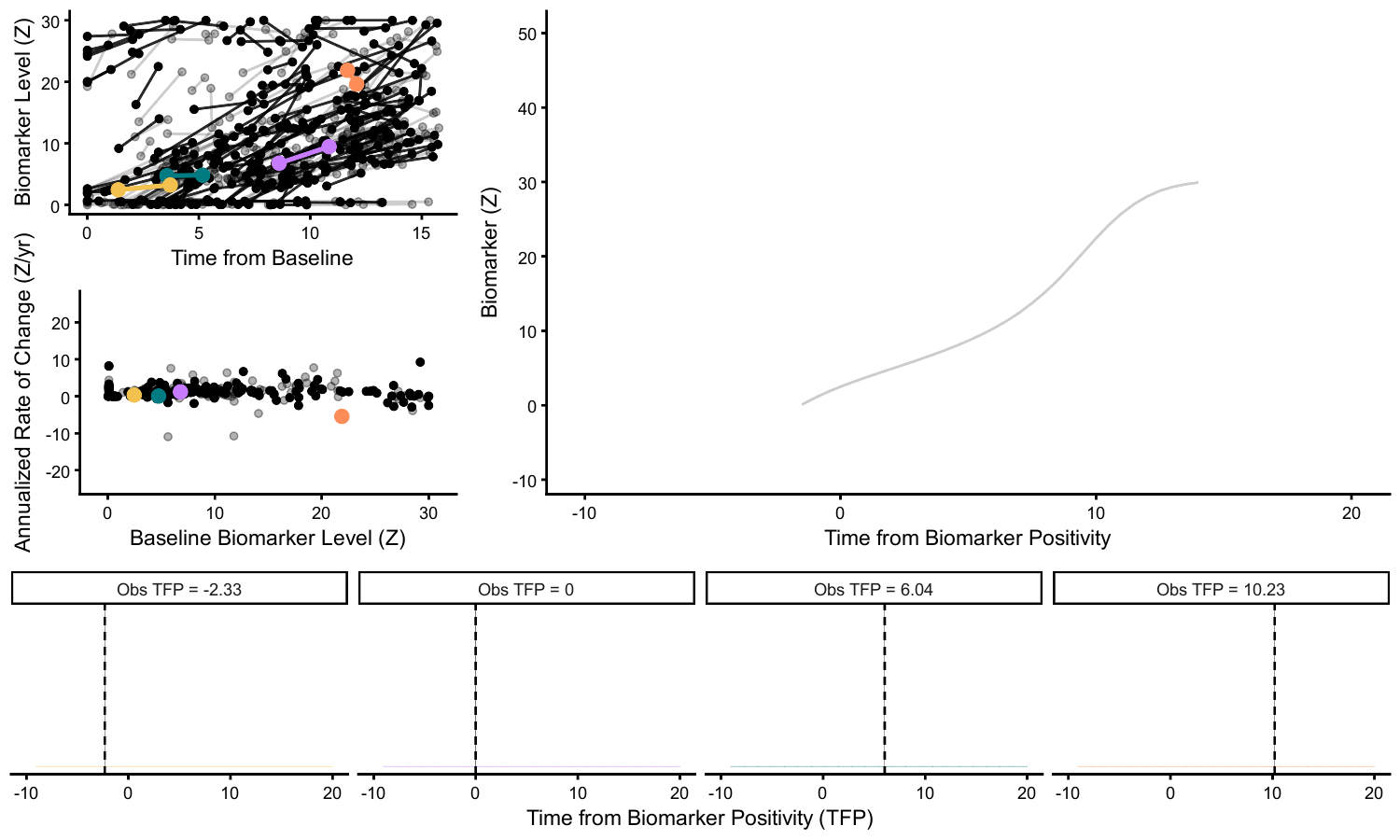

### Supp Video - DIAN CSF pT181

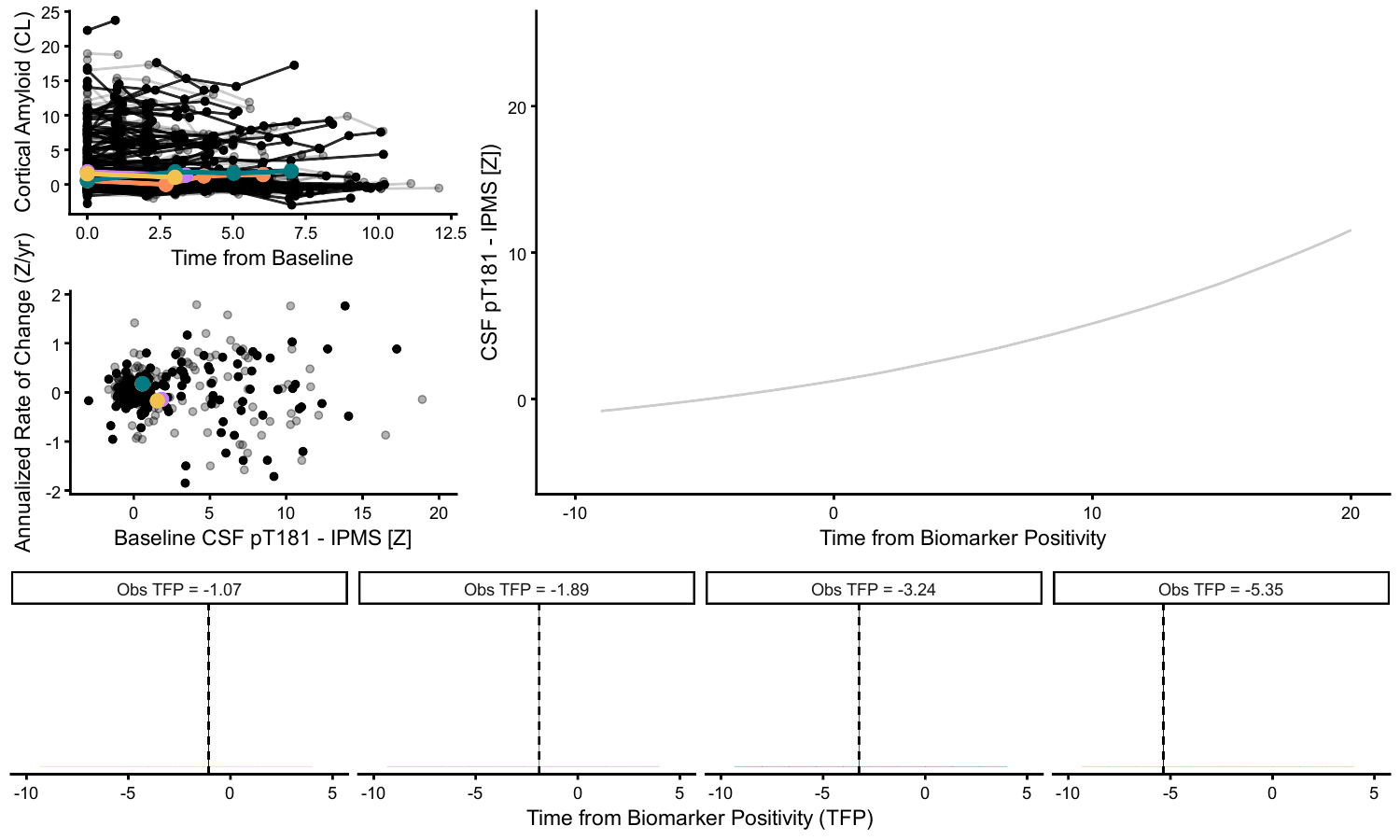

### Supp Video - DIAN PET PiB

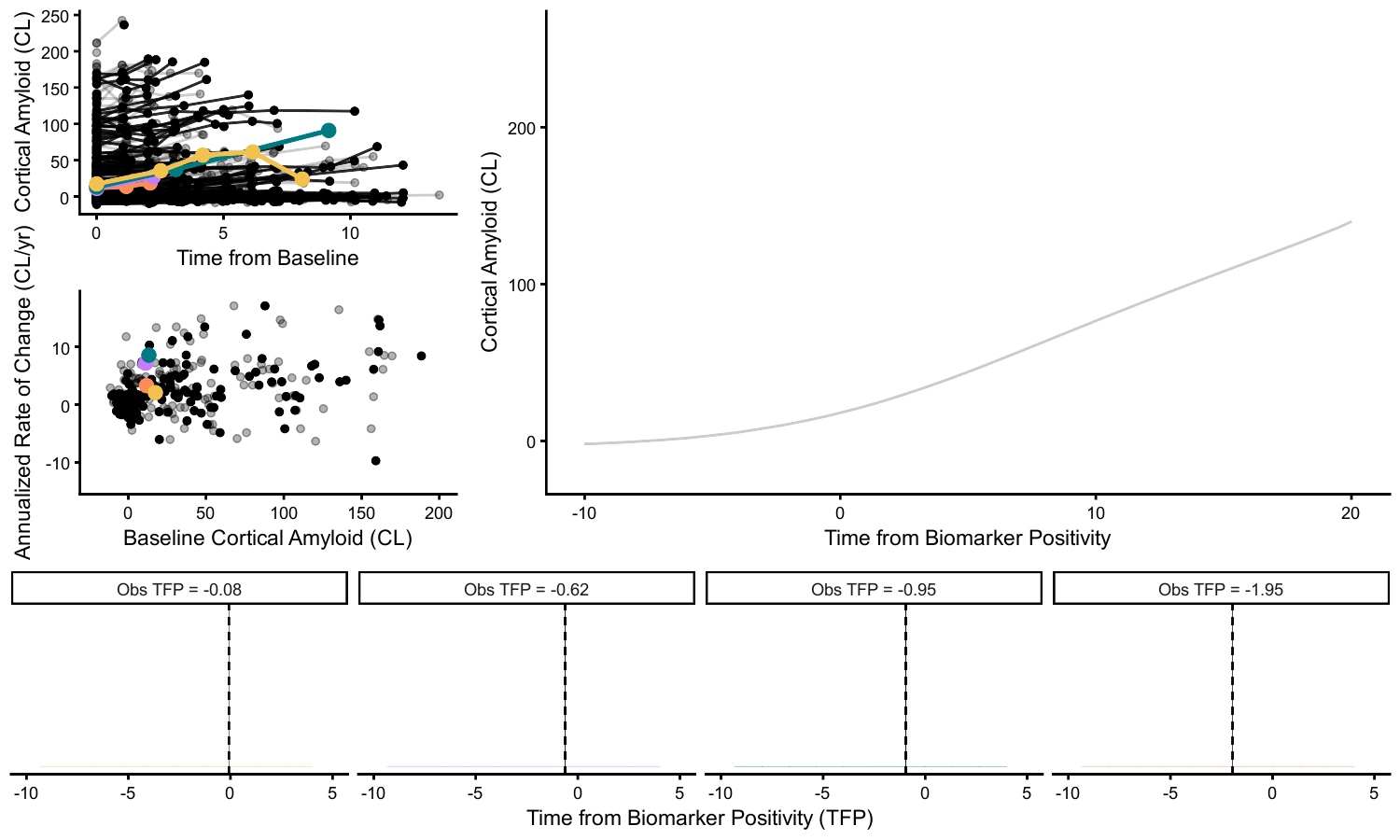
